# An Integrated Single-Nucleus Atlas Resolves Cell-Type-Specific Programs and Molecular Subtypes in Alzheimer’s Disease

**DOI:** 10.64898/2026.09.03.747935

**Authors:** Negin Rahimzadeh, Samuel Morabito, Saniya Khullar, Zechuan Shi, Zhenkun Cao, Vivek Swarup

## Abstract

Interindividual heterogeneity in Alzheimer’s disease (AD) remains poorly understood, as disparate single-cell studies leave it unclear whether findings reflect shared architecture or dataset-specific idiosyncrasies. Here, we present panAD, a transcriptomic atlas of >3 million nuclei from 791 individuals across 13 studies, spanning AD, mild cognitive impairment, and cognitively normal aging. AD converges on a reproducible, cell-type-specific molecular architecture: co-expression modules track neuropathology and cognitive decline; GWAS risk genes act predominantly as downstream targets of transcription factor hubs such as microglial *SPI1*; intercellular communication is remodeled with disease stage; and sex differences concentrate in microglial immune-activation programs. To model patient-level transcriptomic heterogeneity, we developed the Multi-seed Optimization of Neural Embeddings for subTyping (MONET) framework, in which a masked variational autoencoder applied to covariate-adjusted, multi-cell-type profiles resolves four subtypes (Metal-Ion Stress, Neuroinflammatory, Synaptic Integrity, and Tissue Remodeling) that dissociate neuropathological burden from cognitive impairment and nominate predominantly non-overlapping candidate therapeutics. Finally, Stellar Atlas provides an AI-native conversational interface to the atlas.

## Introduction

Alzheimer’s disease (AD) is the most prevalent cause of dementia worldwide, with approximately 55 million individuals currently living with dementia, a number projected to triple by 2050 as global populations age^1^. Despite decades of research, disease-modifying therapies remain scarce, partly because the cellular and molecular heterogeneity of AD has not been sufficiently resolved to guide therapeutic development. The emergence of single-nucleus RNA sequencing (snRNA-seq) has created an opportunity to dissect the transcriptional landscape of the human brain at single-cell resolution, enabling characterization of diverse neuronal and non-neuronal populations within a complex, post-mortem tissue environment.

Rather than a single, uniform disease process, Alzheimer’s disease (AD) is increasingly recognized as a heterogeneous disorder that varies across individuals, brain regions, and disease stages^2, 3^. Cellular states differ substantially by sex, genetic background, anatomical location, and neuropathological progression, while selective neuronal vulnerability follows distinct spatial and temporal patterns across the brain^4^. Even the clinical presentation of AD spans a broad spectrum, from rare autosomal-dominant forms with early onset to the much more common late-onset disease. This biological diversity is mirrored by marked regional differences in pathological spread, transcriptional dysregulation, and cellular responses to disease.

Single-cell and single-nucleus transcriptomic studies have transformed our understanding of this heterogeneity by identifying disease-associated microglial states, reactive astrocytes, oligodendrocyte dysfunction, neuronal vulnerability, and numerous cell-type-specific transcriptional programs^5–11^. However, because these studies profile distinct cohorts, brain regions, disease stages, sequencing platforms, and analytical pipelines, it remains unclear whether the cellular states and molecular programs they report converge on a reproducible, shared molecular architecture of AD or instead reflect context- or dataset-specific variation. Resolving this question requires harmonizing diverse datasets within a unified analytical framework rather than interpreting each study in isolation.

The challenge, however, extends beyond increasing sample size. Meaningful integration aims to preserve genuine biological variation while distinguishing reproducible disease-associated features from cohort-specific observations. Differences in clinical and neuropathological metadata, ancestry representation, tissue sampling, sequencing technologies, quality-control procedures, and cell-type annotation strategies remain substantial obstacles to cross-study integration. Consequently, although the field has become increasingly successful at describing average disease-associated transcriptional changes, it remains far less equipped to determine whether these changes differ systematically across patient populations, whether conserved gene regulatory and co-expression programs represent a more reproducible unit of disease biology than individual genes, or whether AD comprises discrete molecular subtypes rather than a continuum of patient-specific transcriptional variation. Moreover, the regulatory programs underlying cell-type-specific transcriptional responses, the intercellular communication networks coordinating disease progression, and the existence of reproducible molecular AD subtypes remain incompletely understood.

Integrating publicly available single-cell datasets within a unified analytical framework directly addresses these challenges. Rather than simply creating a larger atlas, harmonized integration identifies molecular features that consistently replicate across different studies while preserving biologically meaningful heterogeneity, thereby defining the shared molecular architecture of AD. Such a panAD atlas enables reconstruction of cell-type-specific regulatory networks, inference of multicellular communication across disease progression, identification of conserved co-expression programs, and discovery of molecularly defined AD subtypes that would be difficult or impossible to resolve within individual cohorts. In this way, integration transforms disparate transcriptomic studies into a coherent reference framework for investigating the cellular organization of AD across diverse patient populations.

Beyond improving our understanding of disease biology, integrated atlases also provide a foundation for precision medicine. Reproducible molecular subtypes can be linked to perturbational resources such as CMap/CLUE^12^ to prioritize compounds predicted to reverse subtype-specific transcriptional programs, shifting therapeutic discovery from targeting an averaged AD signature toward matching interventions to distinct molecular disease states. As additional cohorts, including genetically defined AD, Down syndrome-associated AD, cognitively resilient individuals, and the oldest-old, are incorporated into harmonized reference atlases, these resources will increasingly capture the full spectrum of human disease. Realizing this vision will require continued efforts toward standardized metadata, harmonized cell-type annotation, broader ancestral representation, and rigorous cross-cohort integration, ensuring that future molecular insights accurately reflect the diversity and complexity of Alzheimer’s disease.

Here, we assembled a comprehensive integrated single-nucleus atlas of the human cortex comprising more than 3 million quality-controlled nuclei from 791 individuals across 13 separate snRNA-seq datasets, applying uniform alignment and analysis protocols. Donors span three diagnostic categories: AD, MCI, and cognitively normal controls. We applied a multi-layer analytical framework integrating weighted gene co-expression network analysis, cell-cell communication inference, gene regulatory network reconstruction, sex-stratified transcriptomic analyses, and deep generative modeling for molecular subtyping (MONET). Through this integrative approach, we delineate cell-type-specific transcriptional modules, intercellular signaling programs, regulatory networks, sex differences, and four molecularly distinct AD subtypes, along with associated drug-repurposing candidates. Collectively, this work provides a mechanistically grounded resource for the AD research community and identifies new directions for precision intervention. Two features distinguish this resource. First, by integrating rather than merely cataloguing these datasets, we define a unified, cross-cohort molecular architecture of AD, a shared set of cell-type-specific programs, regulatory networks and communication changes reproducible across 13 studies. Second, we extend beyond what any single study has done, both analytically, by resolving molecular subtypes of AD, and as a community resource, by releasing an AI-native, chat-based interface (Stellar Atlas) for interrogating the atlas.

Our findings reveal that AD is not a monolithic transcriptional state but a spectrum of molecular programs that differ across cell types, sexes, and disease subtypes. The identification of prodromal communication signatures in MCI, the delineation of sex-biased regulatory programs, and the nomination of subtype-specific drug candidates collectively illustrate the value of large-scale, multi-modal single-cell atlases for advancing mechanistic understanding and therapeutic targeting in neurodegeneration.

## Results

### A harmonized cross-cohort single-nucleus atlas of the aging and AD human cortex

To enable systematic characterization of cell-type-specific and inter-individual transcriptional variation in Alzheimer’s disease (AD), we assembled a harmonized single-nucleus RNA-sequencing (snRNA-seq) atlas spanning 13 published human cortical datasets ^5–8, 10, 13–20^(n = 791 individuals; >3 million nuclei) (Fig. 1a). After quality control, batch integration, and cell-type annotation (Methods), this yielded a unified, cross-cohort atlas of the aging and AD human cortex (the “panAD atlas”). This resource supported two complementary downstream analytic strategies: (i) atlas-wide characterization of cell-type-resolved biological programs altered in AD, including condition- and sex-specific differential expression, cell-type-specific co-expression programs, intercellular communication changes, and gene regulatory network reconstruction; and (ii) an orthogonal, multi-cell-type molecular subtyping framework (MONET; Methods), in which covariate-adjusted, cell-type-specific pseudobulk profiles were used to train a masked variational autoencoder and consensus-cluster individuals into transcriptionally and clinically distinct AD subtypes, ultimately nominating subtype-specific candidate treatments.

**Fig. 1.**
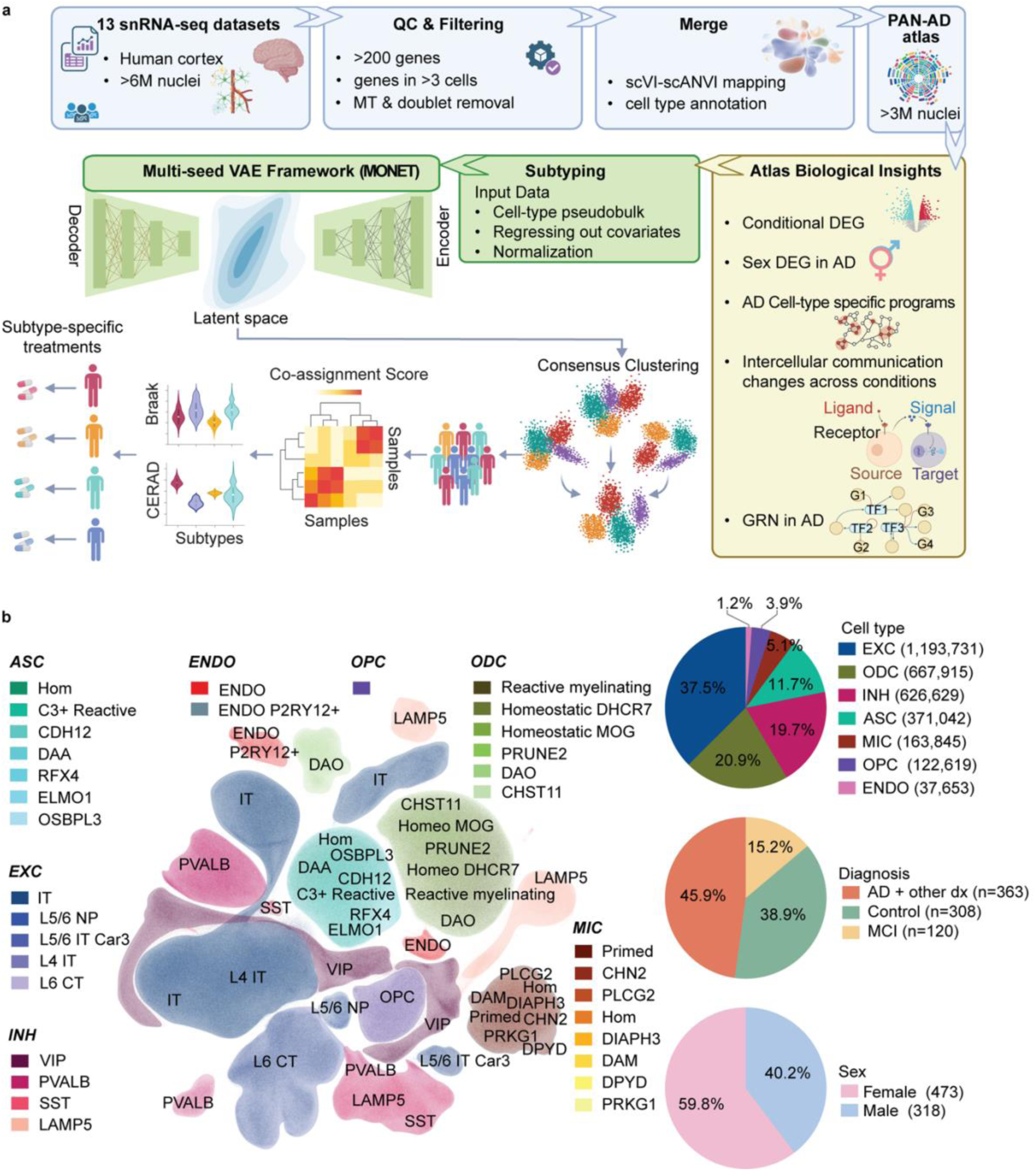
Construction and cellular composition of the panAD single-nucleus transcriptomic atlas. **a,** Schematic of atlas construction and the overall analytical workflow. Raw single-nucleus RNA-sequencing data from 13 published human cortical datasets (n = 791 donors) were quality-filtered (>200 genes per nucleus, genes detected in >3 nuclei, removal of nuclei with high mitochondrial-read content or doublet calls) and integrated across cohorts using scVI-scANVI pipelines, followed by cell-type annotation to generate the harmonized panAD atlas. Downstream analyses branch into (i) atlas characterization and biological-insight analyses, comprising diagnosis-conditional differential gene expression (DEG), sex-stratified DEG in AD, cell-type-specific co-expression programs, intercellular communication changes across diagnostic groups, and gene regulatory network (GRN) inference, and (ii) an unsupervised subtyping pipeline in which cell-type pseudobulk profiles are covariate-regressed and normalized, used to train a multi-seed masked variational autoencoder (encoder-latent space-decoder), and clustered by consensus clustering (seed co-assignment heatmap) to define four reproducible AD molecular subtypes (AD1-AD4) that differ in neuropathological burden (Braak stage, CERAD score) and candidate subtype-specific therapeutic compounds. **b,** UMAP embedding of all profiled nuclei colored by fine-grained cell substate (33 substates spanning EXC, INH, ODC, ASC, MIC, OPC, and ENDO lineages), with the proportion and number of nuclei contributed by each substate indicated in the legend. Top pie chart summarizes atlas nuclei composition by major cell types. The middle and bottom pie charts show donor counts. EXC, excitatory neurons; INH, inhibitory neurons; MIC, microglia; ASC, astrocytes; ODC, oligodendrocytes; OPC, oligodendrocyte progenitor cells; END, endothelial cell. Dx, diagnosis.

### Cell-type and disease-state composition of the panAD atlas

Cell-type annotation of the merged atlas identified seven major cortical cell classes; excitatory neurons, 37.5%, n =1,193,731; oligodendrocytes, 20.9%, n = 667,915; inhibitory neurons, 19.7%, n = 626,629; astrocytes, 11.7%, n = 371,042; microglia, 5.1%, n = 163,845; OPCs, 3.9%, n = 122,619; Endothelial cells, 1.2%, n=37,653 (Fig. 1b). Astrocytes and microglial populations were further resolved into homeostatic and disease/reactive substates.

Glial populations in particular captured well-described disease-associated states, including reactive/myelinating oligodendrocytes, C3-reactive and disease-associated astrocytes (ASC_C3_Reactive and ASC_DAA), and microglial states including primed (MIC_Primed) and a small population of disease-associated microglia (MIC_DAM), alongside their respective homeostatic counterparts (ODC_DHCR7_Hom, ASC_Hom, MIC_Hom) (Fig. 1b). Excitatory and inhibitory neurons were further resolved into canonical cortical subtypes (e.g., EXC_IT, EXC_L5/6 NP, EXC_L4 IT, EXC_L6 CT; INH_VIP, INH_PVALB, INH_SST, INH_LAMP5), recapitulating expected cortical neuronal diversity. The relatively small MIC_DAM population (n = 760) is consistent with previous reports that canonical disease-associated microglia are comparatively heterogeneous and less discretely represented in human cortical tissue than in mouse models^21^ (Fig. 1b).

The atlas encompasses AD, control, and MCI groups, with AD accounting for 45.7% (n = 1,453,622), control for 41.7% (n = 1,329,566), and MCI for 12.6% (n = 400,246) of nuclei. Most cells came from female donors with 60.6% (n = 1,928,120), while cells from males were about 39.4% (n = 1,255,314) (Fig. 1b). This sex imbalance motivated explicit covariate adjustment for sex during subtyping and dedicated downstream analysis of sex-specific transcriptional differences in AD, underscoring the importance of accounting for cohort composition when interpreting disease-associated signatures derived from this resource.

### Cell-type-specific co-expression modules in neuronal and glial populations

To characterize transcriptional organization programs within each major cell population, we applied hdWGCNA^22^ (high-dimensional weighted gene co-expression network analysis, a framework we developed for constructing co-expression networks in single-nucleus data) to excitatory and inhibitory neurons, microglia (MIC), oligodendrocytes (ODC), and astrocytes (ASC), constructing weighted gene co-expression networks from metacell-aggregated expression profiles (Figs. 2-3; Methods). This analysis identified 6 excitatory and 12 inhibitory neuron co-expression modules, each with distinct functional enrichment profiles (Fig. 2a). Supplementary Tables 2 and 3 provide cell-type-specific module characterization and module-associated hub genes, respectively.

**Fig. 2.**
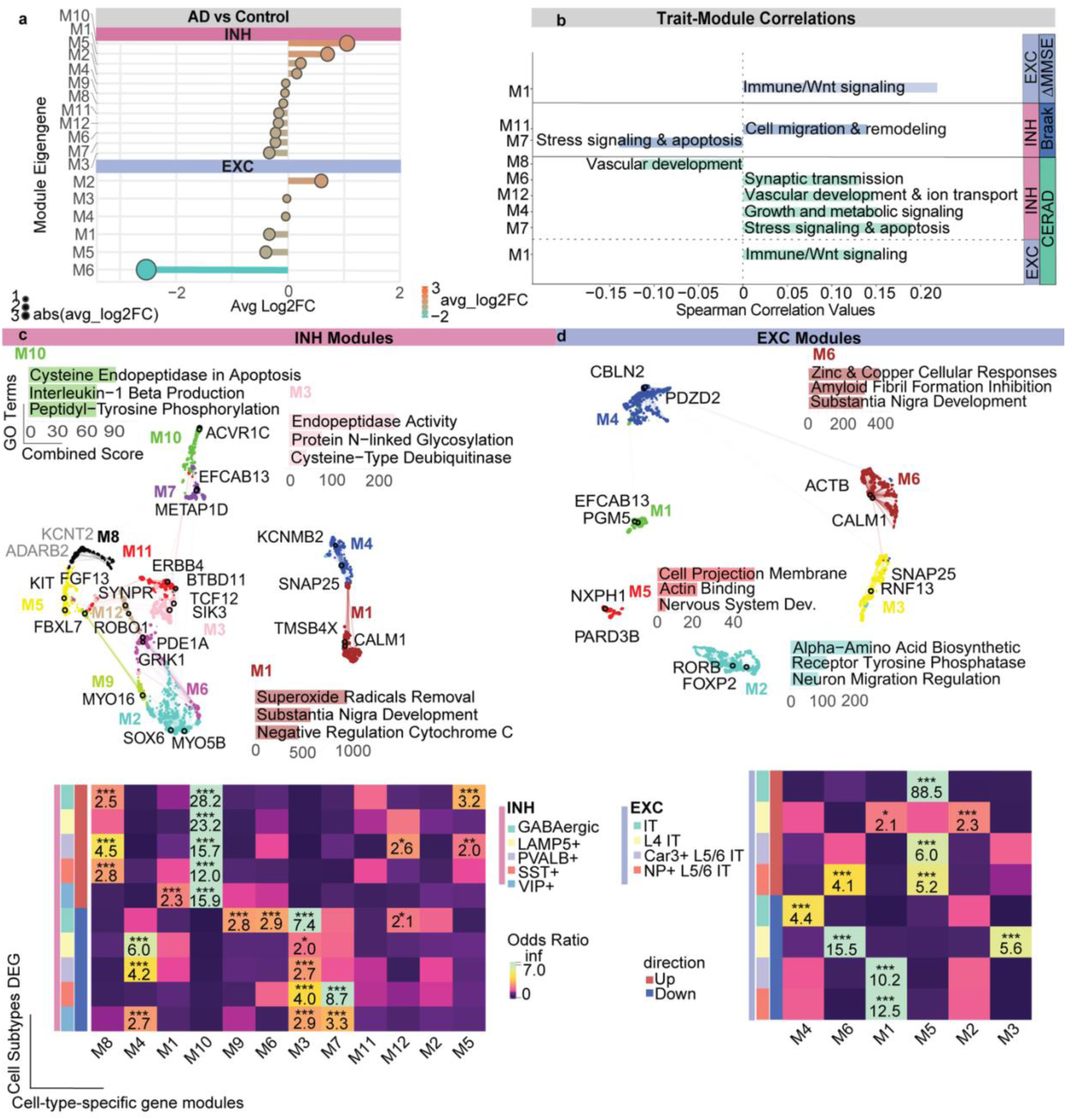
Cell-type-specific co-expression modules reveal divergent transcriptional programs in excitatory and inhibitory neurons in Alzheimer’s disease. **a**, Average log2 fold change (AD versus control) of module eigengene expression for inhibitory (INH, top) and excitatory (EXC, bottom) neuronal co-expression modules identified by high-dimensional weighted gene co-expression network analysis (hdWGCNA). Dot size denotes |avg. log2FC|; dot color denotes signed avg. log2FC. **b**, Spearman correlations between module eigengenes and neuropathological/cognitive traits (Braak stage, ΔMMSE, CERAD score) for INH and EXC modules, annotated with representative enriched biological functions for each trait-associated module (e.g., immune/Wnt signaling, stress signaling and apoptosis, cell migration and remodeling, vascular development, synaptic transmission, growth and metabolic signaling). Module-trait correlations computed in ROSMAP samples only (n=176). Bars show significant trait-module Spearman correlations (FDR < 0.05). **c**, Top enriched Gene Ontology (GO) terms (ranked by combined score) and hub-gene co-expression networks for INH modules M1-M12 (e.g., M10: cysteine endopeptidase activity in apoptosis, interleukin-1β production, peptidyl-tyrosine phosphorylation; M1: superoxide radical removal, substantia nigra development, negative regulation of cytochrome c; M3: endopeptidase activity, protein N-linked glycosylation, cysteine-type deubiquitinase activity), with representative hub genes labeled (e.g., *ACVR1C*, *EFCAB13*, *METAP1D*, *KIT*, *FBXL7*, *MYO5B*, *SOX6*, *SNAP25*, *KCNMB2*, *TMSB4X*, *CALM1*). **d**, Top enriched GO terms and hub-gene co-expression networks for EXC modules M1-M6 (e.g., M6: zinc and copper cellular responses, amyloid fibril formation inhibition, substantia nigra development; M5: cell projection membrane, actin binding, nervous system development; M2: alpha-amino acid biosynthetic process, receptor tyrosine phosphatase activity, neuron migration regulation), with representative hub genes labeled (e.g., *CBLN2*, *PDZD2*, *ACTB*, *CALM1*, *SNAP25*, *RNF13*, *NXPH1*, *PARD3B*, *RORB*, *FOXP2*). Heatmaps below panels c and d show odds ratios for the enrichment of cell-subtype-specific up- and downregulated DEGs (INH subtypes: GABAergic, LAMP5+, PVALB+, SST+, VIP+; EXC subtypes: IT, L4 IT, Car3+ L5/6 IT, NP+ L5/6 IT) within each co-expression module (***FDR < 0.001; color denotes odds ratio; red/blue side bars denote DEG direction). Annotated heatmap cells indicate comparisons with FDR < 0.05, q ≥ 4, and overlap ≥ 2%.

**Fig. 3.**
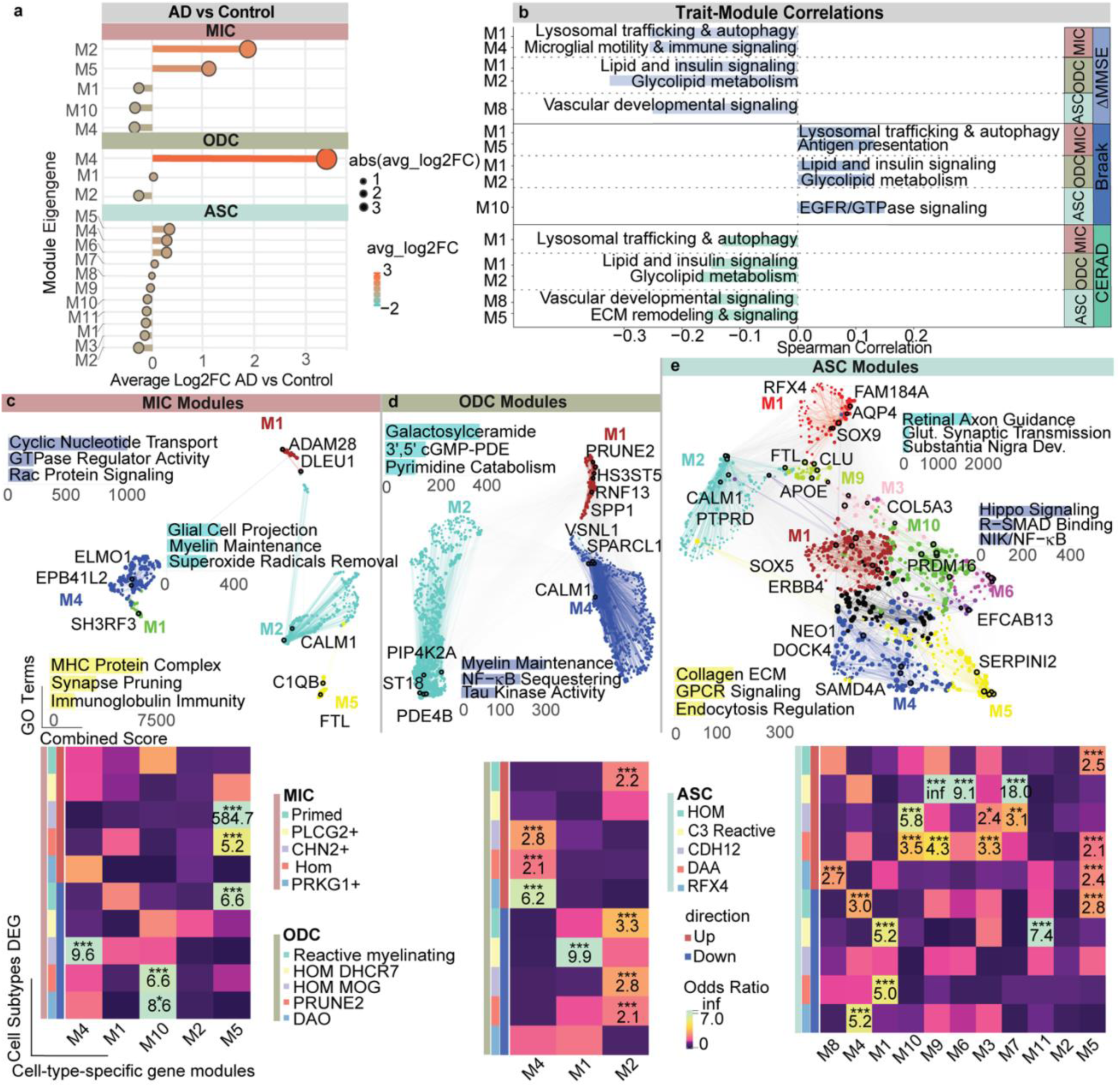
Cell-type-specific co-expression modules reveal divergent transcriptional programs in microglia, oligodendrocytes, and astrocytes in Alzheimer’s disease. **a**, Average log2 fold change (AD versus control) of module eigengene expression for microglial (MIC, top), oligodendrocyte (ODC, middle), and astrocyte (ASC, bottom) co-expression modules identified by hdWGCNA. Dot size denotes |avg. log2FC|; dot color denotes signed avg. log2FC. **b**, Spearman correlations between module eigengenes and neuropathological/cognitive traits (Braak stage, ΔMMSE, CERAD score) for MIC, ODC, and ASC modules, annotated with representative enriched biological functions (e.g., lysosomal trafficking and autophagy, microglial motility and immune signaling, antigen presentation, lipid and insulin signaling, glycolipid metabolism, EGFR/GTPase signaling, vascular developmental signaling, ECM remodeling and signaling). Module-trait correlations computed in ROSMAP samples only. Bars show significant trait-module Spearman correlations (FDR < 0.05). **c**, Top enriched GO terms and hub-gene co-expression networks for MIC modules (e.g., glial cell projection, myelin maintenance, superoxide radical removal; M5: MHC protein complex, synapse pruning, immunoglobulin-mediated immunity), with representative hub genes labeled (e.g., *ELMO1*, *EPB41L2*, *SH3RF3*, *ADAM28*, *DLEU1*, *CALM1*, *C1QB*, *FTL*). **d**, Top enriched GO terms and hub-gene co-expression networks for ODC modules (e.g., galactosylceramide metabolism; cyclic GMP-phosphodiesterase activity and pyrimidine catabolism; myelin maintenance, NF-κB sequestering, tau kinase activity), with representative hub genes labeled (e.g., *PRUNE2*, *HS3ST5*, *RNF13*, *SPP1*, *VSNL1*, *SPARCL1*, *PIP4K2A*, *ST18*, *PDE4B*, *FTL*). e, Top enriched GO terms and hub-gene co-expression networks for ASC modules (e.g., collagen/ECM organization, GPCR signaling, *SAMD4A*-mediated endocytosis regulation; Hippo signaling, R-SMAD binding, NF-κB sequestering), with representative hub genes labeled (e.g., *RFX4*, *FAM184A1*, *AQP4*, *CLU*, *SOX9*, *APOE*, *PTPRD*, *CALM1*, *COL5A3*, *ERBB4*, *SOX5*, *NEO1*, *DOCK4*, *EFCAB13*, *SERPINI2*). Heatmaps below panels c-e show odds ratios for the enrichment of cell-subtype-specific up- and downregulated DEGs (MIC subtypes: Primed, PLCG2+, CHN2+, Hom, PRKG1+; ODC subtypes: Reactive myelinating, HOM DHCR7, HOM MOG, PRUNE2+, DAO; ASC subtypes: Hom, C3+ Reactive, CDH12+, DAA, RFX4+) within each co-expression module (***FDR < 0.001; color denotes odds ratio; red/blue side bars denote DEG direction). Annotated heatmap cells indicate comparisons with FDR < 0.05, q ≥ 4, and overlap ≥ 2%.

Differential module eigengene (DME) analysis comparing AD to cognitively normal controls demonstrated significant AD-associated upregulation and downregulation of specific modules in both cell types, implicating transcriptional reprogramming of synaptic, metabolic, and stress-response programs as integral components of neuronal vulnerability in AD (Fig. 2a).

Multiple modules showed significant correlations with Braak stage, CERAD score, and longitudinal cognitive decline (ΔMMSE) (all FDR < 0.05; Fig. 2b). In inhibitory neurons, stress signaling and apoptosis were represented by INH-M7, which was downregulated in AD yet significantly associated with Braak and CERAD staging scores (Fig. 2a, b).

Inflammatory signaling was a prominent feature of INH-M10, the most upregulated inhibitory neuronal module in AD (Fig. 2c). Oxidative stress, metal-ion homeostasis, and cytoskeletal stability characterized INH-M1, the second most upregulated INH module in AD with the hub genes *CALM1* and *TMSB4X*. INH-M1 was also significantly enriched in VIP interneurons (odds ratio = 2.3) (Fig. 2c).

Within the excitatory neuronal populations, neuronal identity and axon-guidance programs were upregulated in AD, represented by M2, characterized by *RORB* and *FOXP2* (Fig. 2a, d). In contrast, excitatory programs encompassing *NXPH1* and *PARD3B* (M5), and *ACTB* and *CALM1* (M6) were downregulated in AD. M5 was associated with intracellular signaling, whereas M6 was enriched for zinc- and copper-ion homeostasis with inhibition of amyloid fibril formation (Fig. 2a, d). Immune-related signaling (including Wnt) was additionally linked to clinical and pathological features, with EXC-M1 representing the only excitatory module significantly associated with CERAD and ΔMMSE (Fig. 2b). GO enrichment analyses indicated that M1 was involved in signaling and immune pathways; although ambient RNA was reduced, this immune enrichment may reflect neuronal stress-associated immune signaling, residual ambient RNA, or both.

Several glial modules show significant association with Braak stage, CERAD, and ΔMMSE (Fig. 3b). Myelin maintenance and neuronal support characterized ODC-M4, which exhibited the strongest AD-associated transcriptional shifts among oligodendrocyte modules, although it was not significantly associated with clinical or neuropathological traits (Fig. 3a, b). In contrast, lipid metabolism and insulin signaling (ODC-M1) and glycolipid metabolism and membrane remodeling (ODC-M2) were significantly associated with ΔMMSE and neuropathological staging measures (Fig. 3b).

Microglial modules encompassed programs related to immune signaling, lysosomal function, and protein clearance. Lysosomal trafficking and autophagy, represented by MIC-M1, were significantly associated with Braak, CERAD, and ΔMMSE traits. Microglial motility and immune signaling characterized MIC-M4, the most downregulated microglial module in AD, which was also significantly associated with ΔMMSE. Antigen presentation and immune activation characterized MIC-M5, which was upregulated in AD compared to control and significantly associated with Braak staging. Interestingly, neuronal connectivity-related MIC-M2 was the most upregulated microglial module in AD yet showed no significant associations with the examined clinical or neuropathological traits (Fig. 3b). This dissociation suggests that MIC-M2 may represent an early or pathology-independent microglial activation program that is uncoupled from conventional measures of disease severity.

Extracellular matrix (ECM) remodeling and signaling characterized ASC-M5, the most upregulated astrocyte module in AD, which was also significantly correlated with CERAD score. ASC-M10, enriched for EGFR/GTPase signaling, was positively correlated with Braak staging despite only modest changes in overall AD expression (Fig. 3a, b). These astrocytic programs are consistent with a context-dependent shift from homeostatic to reactive astrocyte states. Together, these findings highlight coordinated but cell-type-specific transcriptional adaptations across the major non-neuronal populations of the AD cortex (Fig. 3c-e). Several of these modules re-emerge below as defining features of the four AD molecular subtypes (Fig. 8a).

### Disease-stage-specific remodeling of intercellular communication networks

To characterize how intercellular communication changes across the AD continuum, we inferred cell-cell signaling separately in control, MCI, and AD samples (Fig. 4; Methods). We interpreted differences across control, MCI, and AD as reflecting stage-associated changes in cell-cell signaling, with an overall decline across the disease continuum.

**Fig. 4.**
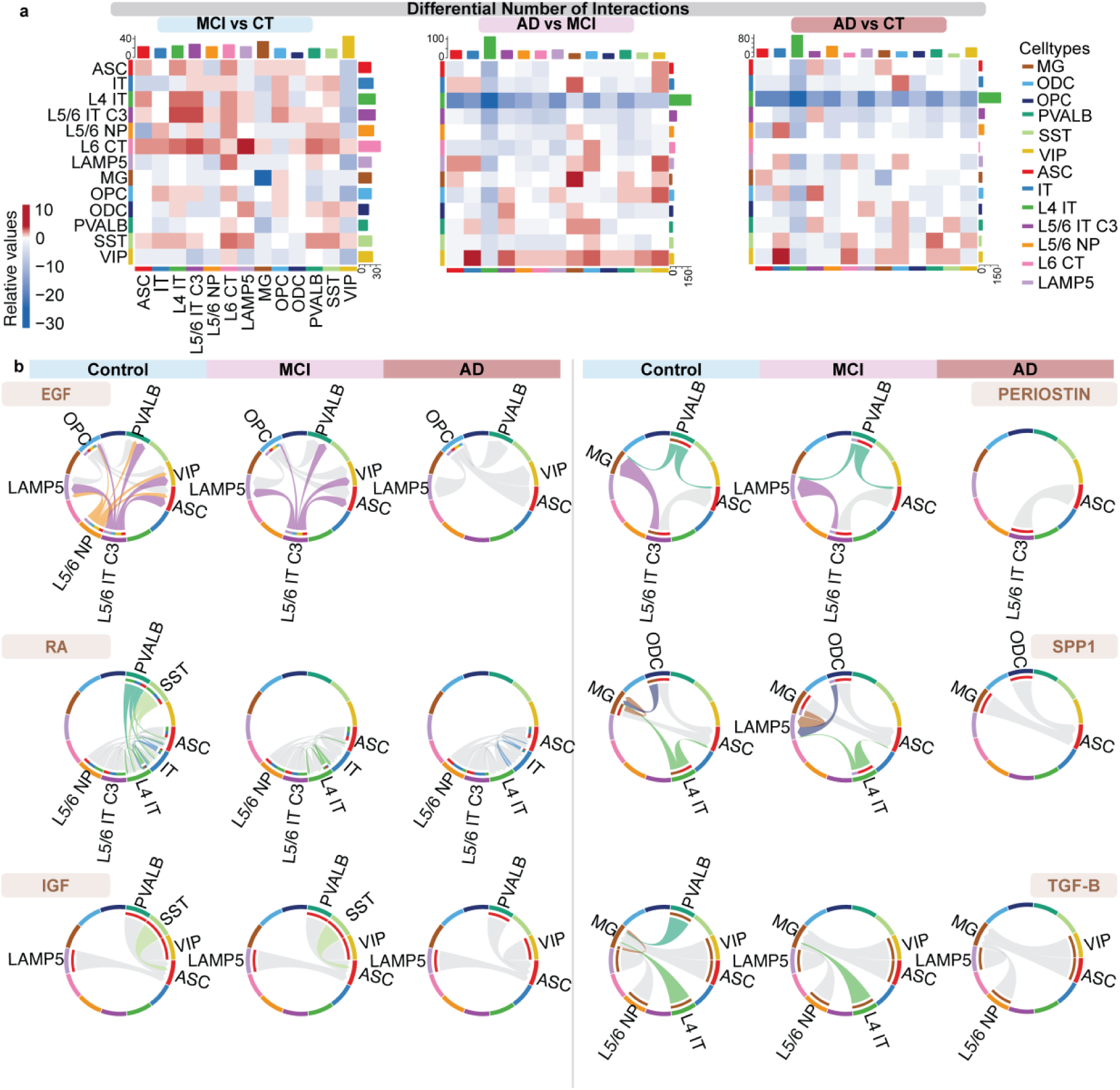
Intercellular communication is progressively lost across multiple signaling pathways in AD. **a**, Heatmaps of the differential number of predicted cell-cell interactions between MCI and Control (left), AD and MCI (middle), and AD and Control (right), across 13 annotated cell types (ASC, IT, L4 IT, L5/6 IT C3, L5/6 NP, L6 CT, LAMP5, MG, OPC, ODC, PVALB, SST, VIP). Heatmap color indicates the relative change in the number of interactions for each sender (row) to receiver (column) cell-type pair (red, increased; blue, decreased, in the later relative to the earlier condition of each comparison). Bar plots flanking each heatmap (top and right) show the total differential number of interactions per cell type, summed across all partner cell types and colored by cell type as indicated in the legend. **b**, Chord diagrams showing predicted cell-cell signaling for six pathways in Control, MCI, and AD. Left: EGF, retinoic acid (RA), and IGF. Right: PERIOSTIN, SPP1, and TGF-β. Arcs represent the indicated cell types; ribbons indicate predicted ligand-receptor signaling between sender and receiver cell types, with ribbon width proportional to signaling strength and gray ribbons indicating signaling absent or below threshold. EGF, IGF, PERIOSTIN, and SPP1 signaling are present in Control and partially maintained in MCI but are lost in AD; RA and TGF-β signaling decline from Control through MCI to AD.

Differential interaction analysis across pairwise comparisons (MCI vs. Control, AD vs. MCI, and AD vs. Control) revealed extensive and coordinated remodeling of cell-cell signaling networks with disease stages (Fig. 4a). Specifically, L4 IT neurons show a loss in the number of differential interactions from control to MCI, and in AD compared to control and MCI (Fig. 4a). Differential number of interactions seems to be lost more in AD vs control, and in AD vs MCI than in MCI compared to control (Fig. 4a). In control brains, EGF signaling was prominently active across multiple sender-receiver cell-type pairs; this pathway was markedly diminished in AD, suggesting an early loss of neuroprotective growth factor signaling as an early feature of disease (Fig. 4b). Several signaling pathways were specifically diminished in AD relative to control. Retinoic Acid (RA) signaling is particularly diminished in PVALB and SST neurons (Fig. 4b). The insulin-like growth factor (IGF) signaling pathway also shows decreased signaling from SST neurons to astrocytes (Fig. 4b). Periostin-mediated communication from PVALB and L5/6 IT C3 neurons to LAMP5 neurons is reduced in AD relative to MCI, while communication from these neuronal populations to microglia is reduced relative to control (Fig. 4b). SPP1 signaling from ODC to microglia and LAMP5 neurons, as well as from Layer 4 IT neurons to LAMP5 neurons and astrocytes are diminished in AD (Fig. 4b). TGF-β signaling from PVALB and L4 IT neurons to microglia is also largely absent in AD (Fig. 4b). This analysis provides a systems-level view of key intercellular signaling being dysregulated as AD progresses from preclinical to symptomatic stages.

### Cell-type-specific gene regulatory networks underlying AD-associated transcriptional programs

Co-expression modules identify genes that vary together (Figs. 2-3). However, because they are symmetric by construction, they cannot distinguish upstream regulators from downstream targets or determine which genes drive a module’s association with disease. To move beyond co-expression to directional regulatory relationships, we integrated pySCENIC with network-constrained regression (NetREm) to infer cell-type-specific gene regulatory networks (GRNs) (Fig. 5). Briefly, pySCENIC defined motif-pruned TF-target (TF-TG) regulons, and NetREm then prioritized interactions supported by both co-expression and protein-protein interaction topology and classified TF-TF relationships as cooperative or antagonistic (Methods)^23–25^.

**Fig. 5.**
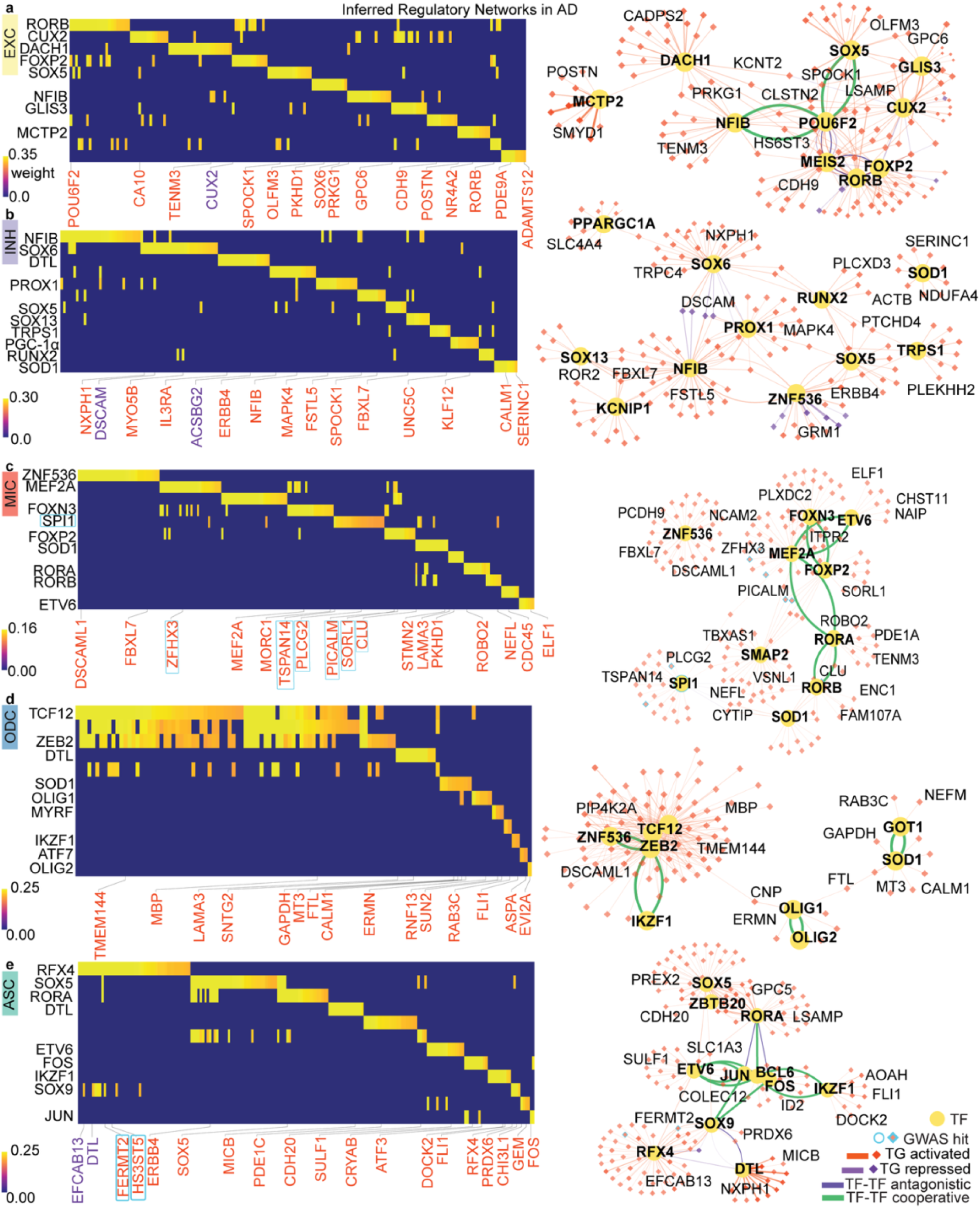
Cell-type-specific gene regulatory networks reveal hub transcription factors and AD GWAS target genes. **a-e,** Inferred regulatory networks for excitatory neurons (EXC; **a**), inhibitory neurons (INH; **b**), microglia (MIC; **c**), oligodendrocytes (ODC; **d**), and astrocytes (ASC; **e**). For each cell type, heatmaps (left) show transcription factors (TFs) with the highest regulon activity and their target genes (TGs), colored by GWAS status (genes with a cyan outline are AD GWAS hits, e.g., *PICALM*, *SORL1*, *CLU*, *PLCG2*, *ZFHX3*, and *TSPAN14* in microglia; *FERMT2* and *HS3ST5* in astrocytes). Heatmap color indicates inferred regulatory edge weight (scale shown per cell type, range 0-0.16 to 0-0.35 depending on cell type). Network graphs (right) depict the corresponding TF-TG and TF-TF regulatory relationships. Yellow circles represent TFs (node size scales with network connectivity; bold labels indicate the most highly connected hub TFs). Red diamonds indicate activated target genes; purple diamonds indicate repressed target genes; light-blue-outlined diamonds indicate AD GWAS hits. Edges indicate TF-TG activation (orange) or repression (purple), and TF-TF cooperative (green) or antagonistic (magenta) relationships, as shown in the legend.

Across all five cell types, the resulting regulons were each anchored by a small set of high-connectivity “hub” TFs rather than being diffusely distributed, and several network modules linked by cooperative (green) TF-TF edges recurred within each cell type, indicating that a limited number of co-acting regulators account for most of the shared regulatory signal.

In excitatory neurons the EXC regulon was dominated by *RORB*, *CUX2*, *DACH1*, *FOXP2*, and *SOX5*, with *NFIB*, *GLIS3*, and MCTP2 contributing smaller target sets (Fig. 5a). In the network view, *DACH1*, *SOX5*, *POU6F2*, *NFIB*, and *CUX2* formed a cooperatively linked hub (green TF-TF edges) regulating a shared neighborhood of targets (e.g., *SPOCK1*, *CLSTN2*, *LSAMP*, *KCNT2*, *HS6ST3*), consistent with a layer-defining transcriptional program acting through a coordinated rather than independent set of regulators.

In inhibitory neurons *NFIB*, *SOX6*, *PROX1*, *SOX5*, *SOX13*, *TRPS1*, PGC-1α (*PPARGC1A*), *RUNX2*, and *SOD1* were the principal TFs (Fig. 5b). *ZNF536* and *NFIB* occupied central positions in the network, bridging multiple TF modules (e.g., *SOX13*/*KCNIP1* and *SOX5*/*TRPS1*) to a common target pool including *DSCAM*, *MAPK4*, *FSTL5*, and *ERBB4*, suggesting *NFIB* and *ZNF536* act as integrating nodes across otherwise distinct INH sub-programs.

In microglia, *SPI1*, *ZNF536*, *MEF2A*, *FOXN3*, *FOXP2*, *ETV6*, *RORA*, and *RORB* formed the MIC regulon (Fig. 5c). Notably, *SPI1* (PU.1), the canonical microglial master regulator and an established AD GWAS risk gene, was linked to multiple downstream AD GWAS target genes, including *PLCG2* and *TSPAN14* (Fig. 5c). These findings indicate that microglial transcription factor programs converge on multiple genetically implicated AD loci, highlighting a shared regulatory architecture connecting genetic risk to downstream transcriptional regulation. A separate cooperative cluster linked *FOXN3*, *ETV6*, *MEF2A*, and *FOXP2*.

In oligodendrocytes, the ODC regulon split into two largely independent modules: a *TCF12*-*ZEB2*-*ZNF536* hub cooperatively linked to *IKZF1* and driving myelin-associated targets (*MBP*, *TMEM144*, *DSCAML1*, *PIP4K2A*), and a second *OLIG1*-*OLIG2*-*SOD1*-*GOT1* module regulating a distinct target set (*FTL*, *GAPDH*, *MT3*, *CALM1*, *CNP*, *ERMN*) (Fig. 5d). This separation suggests two at least partially uncoupled oligodendrocyte transcriptional circuits, one tied to canonical *OLIG1*/*OLIG2* myelination identity and one organized around *TCF12*/*ZEB2*.

In astrocytes, the ASC regulon was the most densely interconnected, with *RFX4*, *SOX5*, *RORA*, *ETV6*, *FOS*, *JUN*, *IKZF1*, *SOX9*, and *BCL6* forming a single large cooperative hub (extensive green TF-TF edges centered on *ETV6*-*JUN*-*BCL6*-*FOS*-*IKZF1*) regulating targets including *GPC5*, *SLC1A3*, *COLEC12*, and *ID2* (Fig. 5e). Within this module, the AD GWAS genes *FERMT2* and *HS3ST5* emerged as direct targets, positioning them downstream of the astrocyte *JUN*/*FOS*/*BCL6*-centered regulatory circuit.

GWAS targets recur as downstream nodes across the regulatory hierarchy. Across all five cell types, AD/FTD GWAS genes (cyan outlines/boxes, Fig. 5) appeared predominantly as TG nodes rather than as TFs, with the clearest enrichment in microglia (*PICALM*, *SORL1*, *CLU*, *PLCG2*, *TSPAN14*, *ZFHX3*) and a secondary enrichment in astrocytes (*FERMT2*, *HS3ST5*). This pattern is consistent with a model in which common AD risk variants act largely downstream of cell-type-defining TF hubs, modulating the expression of disease-relevant effector genes rather than altering core regulatory identity directly, and is concordant with prior reports placing microglial PU.1 (*SPI1*)-driven programs at the center of AD genetic risk^26^.

### Sex-stratified transcriptional and regulatory landscapes in Alzheimer’s disease

To determine whether transcriptional differences between male and female AD brains reflect coordinated changes in specific molecular programs, we tested genes differentially expressed between sexes specifically within AD (“AD-only” sex DEGs) for enrichment in the cell-type-resolved hdWGCNA co-expression modules found earlier (Fig. 6a, b). Module enrichment was strongly cell-type- and direction-dependent. In excitatory neurons, male-upregulated DEGs were selectively enriched in the axon-guidance-associated M4 module. In inhibitory neurons, female-upregulated DEGs were enriched across several modules associated with development and signaling processes, including proteoglycan neurodevelopment (M2), developmental signal transduction (M5), vascular development and ion transport (M12), growth and metabolic signaling (M4), and vascular development and signaling (M8). In astrocytes, male-upregulated DEGs were enriched in modules associated with ECM remodeling and signaling (M5), neuromorphogenesis and Wnt signaling (M1), and EGFR/GTPase (M10).

**Fig. 6.**
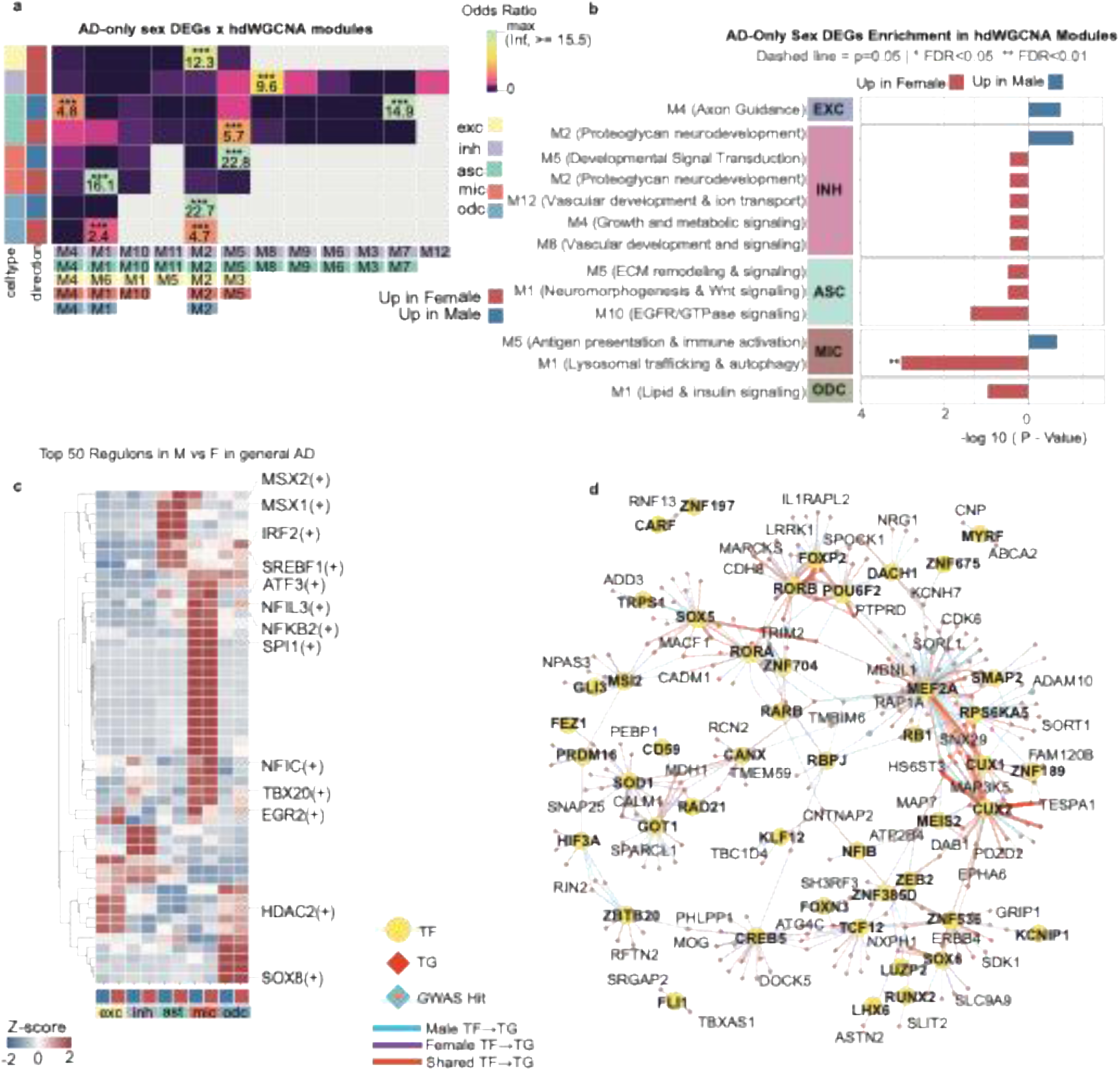
Sex-specific transcriptional and regulatory signatures in AD. **a**, Heatmap of the odds ratio for overlap between AD-only sex-differential genes (DEGs identified between male and female subjects specifically within AD) and hdWGCNA co-expression modules, stratified by cell type (EXC, INH, ASC, MIC, ODC) and direction (up in female, up in male). Color indicates odds ratio (0 to ≥15.5); labeled cells indicate the odds ratio value for significant enrichments (FDR < 0.05, ***). The x-axis shows the cell-type-specific co-expression modules found earlier. **b**, Bar plots of AD-only sex DEG enrichment (−log10 P-value) in hdWGCNA modules, grouped by cell type and colored by direction (up in female, red; up in male, blue). Modules are labeled with their representative biological annotation (e.g., M4, axon guidance; M5, antigen presentation and immune activation; M1, lysosomal trafficking and autophagy). Dashed line indicates P = 0.05; *, FDR < 0.05; **, FDR < 0.01. **c**, Heatmap of the top 50 differentially active regulons between male and female subjects in AD, by cell type. Columns are grouped by cell type (EXC, INH, AST, MIC, ODC) and sex (up in female, up in male, as indicated by the color bar); rows are regulons (transcription factors), hierarchically clustered, with selected regulons labeled (e.g., *MSX2*, *MSX1*, *IRF2*, *SREBF1*, *ATF3*, *NFIL3*, *NFKB2*, *SPI1*, *NFIC*, *TBX20*, *EGR2*, *HDAC2*, *SOX8*). Color indicates row-scaled regulon activity (Z-score, −2 to 2). **d**, Transcription factor-target gene (TF-TG) regulatory network constructed from the top differential regulons shown in c. Yellow circles indicate TFs (node size scales with network connectivity; bolded labels indicate hub TFs); red diamonds indicate target genes (TGs); light-blue diamonds indicate AD GWAS hits among target genes. Edges indicate TF-TG relationships supported predominantly in males (cyan), in females (purple), or shared between both sexes (orange).

The strongest enrichment was observed in microglia. Female-upregulated DEGs were highly enriched in lysosomal trafficking and autophagy program (M1; FDR < 0.01), whereas male-upregulated DEGs were enriched in antigen presentation and immune activation program (M5). Thus, female and male microglia preferentially engaged distinct transcriptional programs in AD, characterized by lysosomal/autophagic and antigen-presentation/immune programs, respectively. These sex-associated microglial programs may contribute to molecular heterogeneity in AD and could influence responses to immunomodulatory therapies.

Oligodendrocyte DEGs upregulated in females were enriched in lipid and insulin signaling represented by M1. Several microglial and neuronal enrichments exhibited odds ratios exceeding 15 (FDR < 0.05; Fig. 6a), indicating that sex-biased transcriptional changes in AD converge on specific, biologically coherent gene programs rather than being diffusely distributed across the transcriptome.

We next asked whether these module-level differences were accompanied by differences in transcription factor (TF) activity. Examining the top 50 regulons differing between male and female AD brains across cell types (Fig. 6c), regulon activity diverged most strongly in glia. A cluster of TFs including *MSX2*, *MSX1*, and *IRF2* showed activity selectively elevated in microglia, while a second cluster, *SREBF1*, *ATF3*, *NFIL3*, *NFKB2*, and *SPI1*, was coordinately upregulated across astrocytes and microglia, consistent with shared activation of lipid-metabolic and immune transcriptional programs in glia. In contrast, *HDAC2* and *SOX8* regulon activity was specifically elevated in oligodendrocytes, consistent with their established roles in oligodendrocyte differentiation and myelination, while *EGR2* activity was enriched in inhibitory neurons. These results suggest that sex differences in AD arise in part from cell-type-restricted reprogramming of specific transcriptional regulators rather than globally altered TF activity. Sex-divergent regulon activity is also concentrated in glial cell types.

To determine whether these sex-biased transcriptional and regulatory programs are accompanied by differences in intercellular communication, we performed sex-stratified CellChat analysis within each diagnostic group (Control, MCI, and AD). Differential interaction number and strength between female and male samples showed distinct cell-type-specific patterns across disease conditions (Supplementary Fig. 2a,b; Supplementary Fig. 3a). Ranked signaling pathway information-flow analysis revealed a pronounced shift in the direction of sex bias in different diseased conditions: in cognitively normal controls and MCI samples, the majority of sex-biased pathways were higher in males, whereas in AD, this pattern reversed, with the majority of sex-biased pathways being higher in females (Supplementary Fig. 3b-d). The majority of the female-biased pathways in AD, including MHC-I, MHC-II, COMPLEMENT, GALECTIN, and UNC5, were male-biased at either both or one of the control or MCI conditions, suggesting that this shift largely reflects reversal of an existing sex bias rather than emergence of a distinct female-specific signaling program (Supplementary Fig. 3b-d).

To identify sex-specific TF-target gene (TG) regulatory systems, we constructed a TF-TG network from the top differential regulons, distinguishing edges supported predominantly in males, in females, or shared between both sexes (Fig. 6d). The network was organized around a small number of highly connected hub TFs, including *MEF2A*, *CUX1*/*CUX2*, *SOX5*, *RORA*/*RORB*, *ZEB2*, and *CREB5*, the majority of whose connections were shared between sexes, suggesting a conserved regulatory backbone. Superimposed on this shared scaffold were sex-specific edges connecting hub and peripheral TFs to distinct sets of target genes. Notably, several network targets correspond to established AD GWAS loci (e.g., *SORL1*, *ADAM10*), indicating that sex-specific regulatory wiring converges on, and may differentially modulate, known AD risk genes. Together, these analyses support a model in which a core, sex-shared regulatory network is differentially modulated in a sex- and cell-type-specific manner in AD, with the most pronounced divergence centered on microglial immune-activation programs. These results show that hub regulators can link sex-specific and shared regulatory networks to AD risk genes.

### A masked variational autoencoder (MONET) identifies four molecularly distinct AD subtypes

To assess molecular heterogeneity among AD individuals beyond cell-type-level analyses, we developed a multi-cell-type subtyping framework integrating covariate-adjusted pseudobulk expression across all seven cell populations (Fig. 7a). We term this framework MONET (Multi-seed Optimization of Neural Embeddings for subTyping). For each sample, the covariate-adjusted pseudobulk profiles of all seven cell types were concatenated into a single multi-cell-type feature vector and encoded by a masked variational autoencoder into a low-dimensional latent representation. Repeating this across 20 random seeds and taking a consensus over the resulting Gaussian-mixture partitions produced four stable subtype assignments (Methods). By modeling all cell types jointly rather than one at a time, MONET captures coordinated cross-cell-type transcriptional variation that per-cell-type analyses cannot resolve. Neff et al. defined three major molecular subtypes of AD using bulk transcriptomics ^27^. To our knowledge, no prior AD single-cell study has performed molecular subtyping at this scale, making this a distinct analytic contribution of the integrated atlas.

**Fig. 7.**
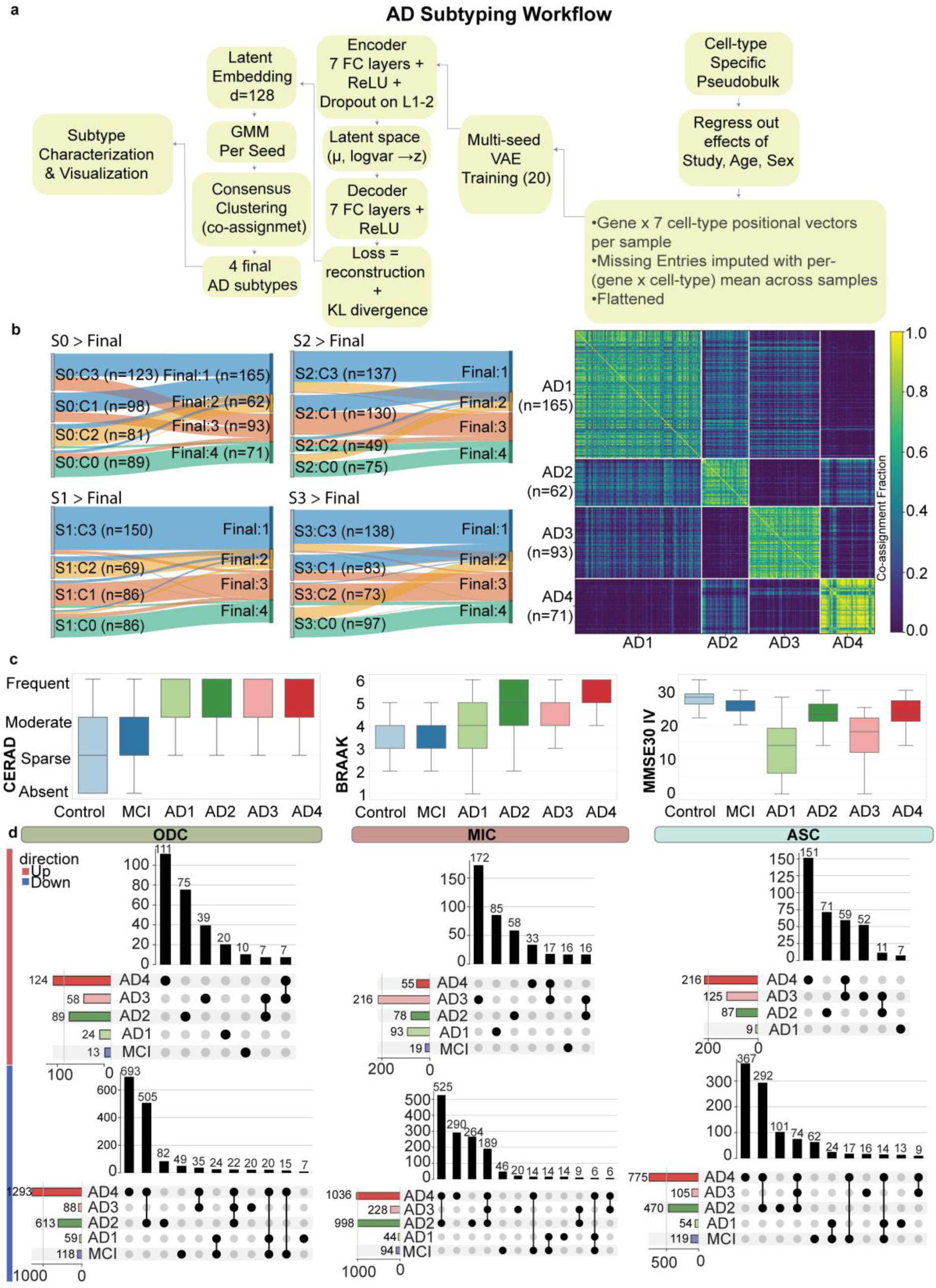
Ensemble VAE-GMM (MONET) identifies four molecularly distinct AD subtypes. **a**, Schematic of the AD subtyping workflow. Cell-type-specific pseudobulk expression profiles were corrected for study, age, and sex effects, concatenated across seven cell types into a fixed-length gene-by-cell-type feature vector per sample (missing entries imputed with the per-gene/cell-type mean), and used to train a masked variational autoencoder (VAE; 20 random seeds, binary masking of imputed/padded positions) to learn a 128-dimensional latent embedding. Gaussian mixture model (GMM) clustering was performed per seed (k = 4, see Methods), and agglomerative hierarchical clustering on the consensus co-assignment matrix across seeds was used to derive four final AD subtypes, which were subsequently characterized and visualized. **b**, Left, Sankey (alluvial) diagrams showing the mapping of cluster assignments from four representative individual seeds (S0-S3; k = 4 each) onto the four final consensus subtypes (Final:1-4; n = 165, 62, 93, and 71 samples, respectively). Right, pairwise sample co-assignment matrix (fraction of seeds in which each pair of samples was assigned to the same cluster), with samples ordered by final consensus cluster membership; block-diagonal structure indicates high clustering stability across seeds. **c**, Distributions of CERAD score (amyloid neuritic plaque density), Braak stage (tau pathology), and MMSE score (global cognition) across Control, MCI, and AD1-AD4 subtypes. Box plots show median, interquartile range (box), and full range (whiskers). CERAD, Braak, and MMSE shown for pooled ROSMAP + SEA-AD samples (n=217); CERAD harmonized across cohorts (see Methods). **d**, UpSet plots showing the number of differentially expressed genes (DEGs, relative to control), split by direction (Up, red; Down, blue) and by group (MCI, AD1-AD4), in oligodendrocytes (ODC), microglia (MIC), and astrocytes (ASC). Horizontal bars indicate total DEG set size per group; vertical bars indicate the size of each intersection, denoted by the connected dots below.

**Fig. 8.**
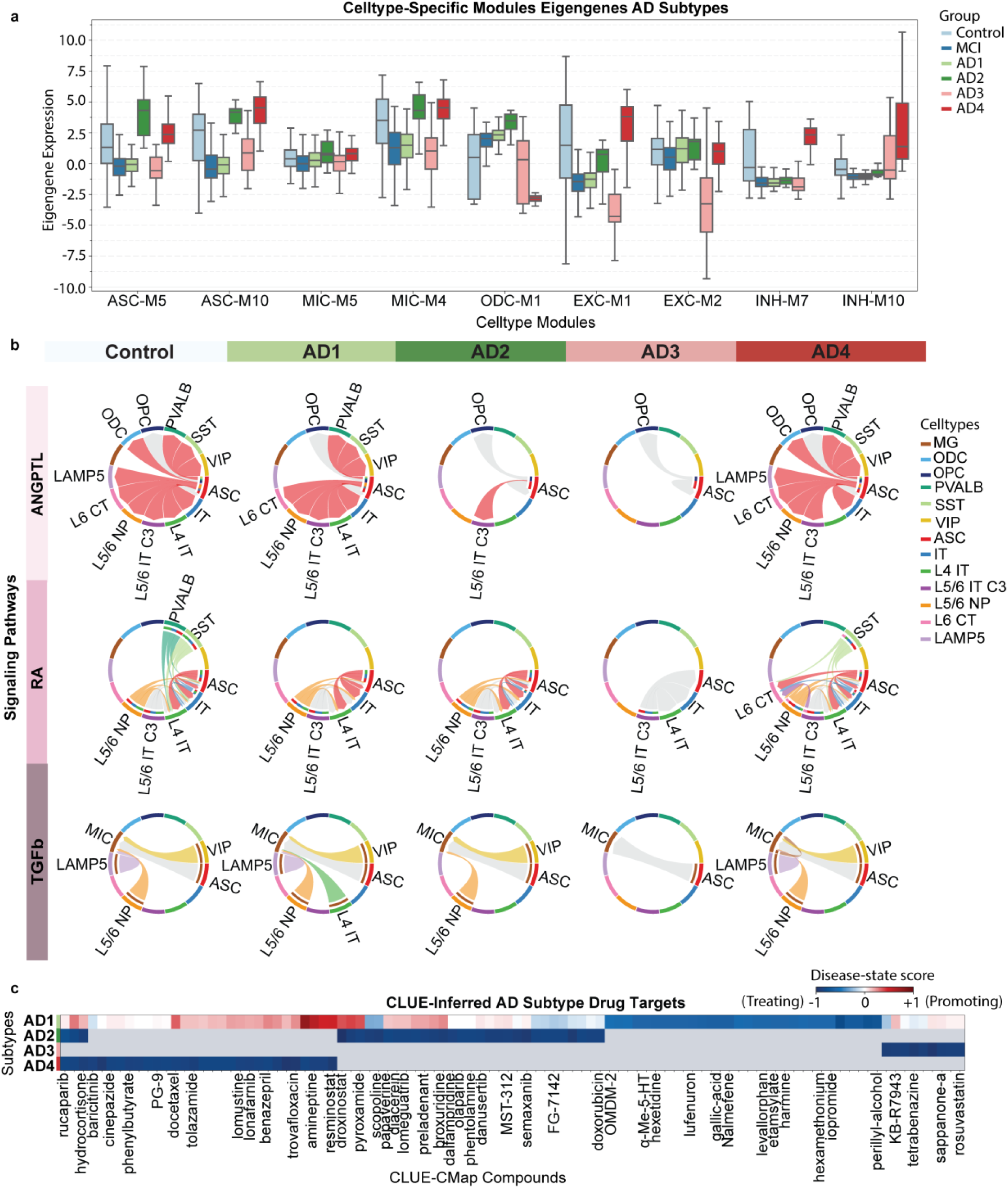
AD molecular subtypes show distinct cell-type co-expression, intercellular signaling, and pharmacological signatures. **a**, Eigengene expression of cell-type-specific co-expression modules (astrocyte, ASC-M5/ASC-M10; microglia, MIC-M5/MIC-M4; oligodendrocyte, ODC-M1; excitatory neuron, EXC-M1/EXC-M2; inhibitory neuron, INH-M7/INH-M10) across Control, MCI, and AD1-AD4 subtypes. Box plots show median, interquartile range, and full range. **b**, Chord diagrams depicting predicted cell-cell signaling for the ANGPTL, retinoic acid (RA), and TGFβ pathways across Control and AD1-AD4 subtypes. Arcs represent annotated cell types (MG, ODC, OPC, PVALB, SST, VIP, ASC, IT, L4 IT, L5/6 IT C3, L5/6 NP, L6 CT, LAMP5; color-coded per legend); ribbons indicate predicted ligand-receptor signaling between cell types, with ribbon width proportional to signaling strength and gray ribbons indicating signaling absent/below threshold. **c**, CLUE/CMap-inferred disease-state scores for candidate compounds, by AD subtype. Heatmap color indicates the similarity of each compound’s transcriptional signature to each subtype’s disease signature (blue, signature-reversing/“treating”; red, signature-mimicking/“promoting”; −1 to +1 scale); gray indicates the compound was not among that subtype’s top-ranked compounds. Colored cells for AD2-AD4 represent compounds meeting the significance threshold; AD1 had none, so its colored cells show its highest-ranked (unfiltered) compounds for comparison only (Supplementary Table 4). Rows: AD subtypes (AD1-AD4); columns: CLUE-CMap compounds.

Across 20 random seeds, silhouette analysis favored a two- or three-cluster solution. We nevertheless selected k = 4 to examine finer-grained molecular heterogeneity, and consensus clustering across seeds yielded AD subtypes (AD1-AD4; n = 165, 62, 93, and 71 samples, respectively) (Supplementary Fig. 4). We refer to these subtypes by their dominant molecular programs, Metal-Ion Stress (AD1), Neuroinflammatory (AD2), Synaptic Integrity (AD3) and Tissue Remodeling (AD4), and use the AD1 to AD4 labels throughout the Results and figures. The block-diagonal structure of the co-assignment matrix, confirmed by pairwise within-versus between-cluster co-assignment distributions, demonstrated high clustering stability across seeds, with individual seed-level cluster assignments (S0-S3) mapping consistently onto the four consensus subtypes (Fig. 7b; Supplementary Fig. 5). To assess whether the AD1-AD4 subtypes reflect biology that is reproducible across another AD cohort and expression platform, we compared subtype-specific transcriptional signatures from our atlas to the independently derived Neff et al. subtypes, defined from bulk transcriptomic profiling of the MSBB cohort. AD1 and AD3 showed robust concordance with two Neff-defined molecular classes (tau/MAPT-predominant and amyloid-predominant/APOE-ε4-enriched, respectively), with more modest concordance observed for AD2 and AD4 (Supplementary Figs. 8 and 9). These results support the external reproducibility of at least a subset of the MONET-derived subtypes across an independent cohort and platform.

**Fig. 9.**
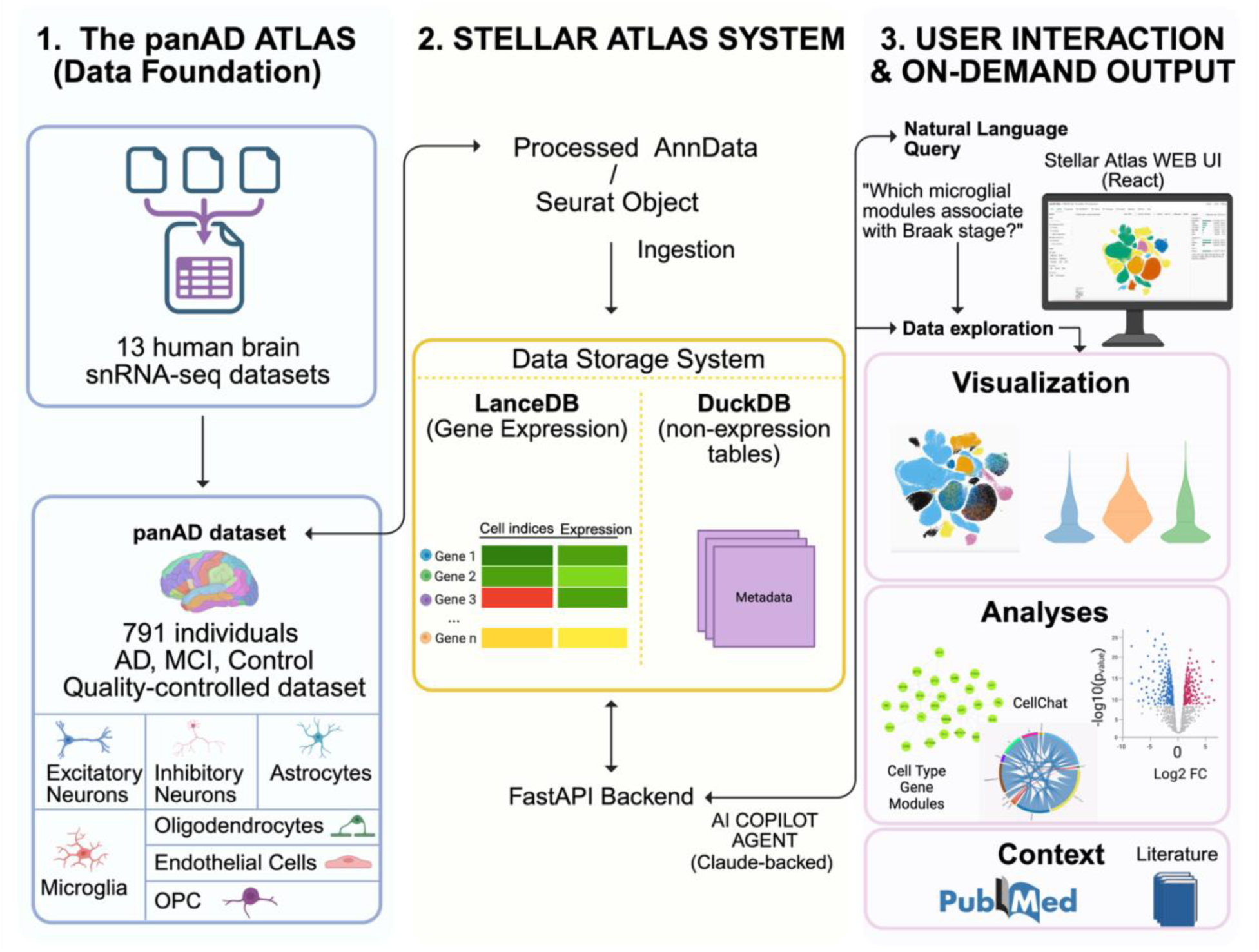
Architecture and functionality of the STELLAR Atlas framework. **a,** Data foundation of the panAD Atlas, comprising 13 harmonized human brain single-nucleus RNA-sequencing (snRNA-seq) datasets from 791 individuals spanning control, mild cognitive impairment (MCI) and Alzheimer’s disease (AD), with major brain cell types represented after quality control. **b,** STELLAR Atlas system architecture. Processed AnnData (.h5ad) or Seurat objects are ingested into complementary data stores: a gene-major LanceDB store for sparse gene-expression data and a DuckDB store for non-expression data, including metadata and precomputed analytical results. A FastAPI backend provides programmatic access to the stored atlas data. A Claude-backed AI copilot enables natural-language interaction with the underlying atlas through tool-based queries. **c,** User interaction and on-demand output. The React-based Stellar Atlas web interface supports interactive data exploration and generation of visualizations and analytical outputs, including gene-expression and UMAP views, differential expression, co-expression modules and cell-cell communication analyses. The copilot enables natural-language queries of the atlas, with optional PubMed retrieval providing literature context for interpretation. This figure was created with BioRender.com.

### Transcriptionally defined AD subtypes differ in the relationship between neuropathological burden and cognitive impairment

We next examined whether the transcriptionally defined AD subtypes differed in neuropathological burden and cognitive status (Fig. 7c). CERAD scores were harmonized onto a common 0-3 scale across cohorts before subtype comparisons (Methods). All AD subtypes exhibited significantly greater neuritic plaque burden (CERAD) and Braak stage than Control and MCI (Fig. 7c). Among the AD subtypes, CERAD scores were broadly similar. In contrast, Braak staging differed more substantially, with AD2 and AD4 showing the highest neurofibrillary tangle burden, whereas AD1 exhibited significantly lower Braak scores than the remaining AD subtypes (Fig. 7c). Cognitive performance varied more markedly than neuropathological measures: AD1 showed the lowest MMSE scores, AD3 displayed intermediate impairment, and AD2 and AD4 retained significantly higher MMSE scores despite similarly high neuropathological burden (Fig. 7c). Together, these findings demonstrate that the relationship between classical neuropathological burden and cognitive impairment differs across transcriptionally defined AD subtypes.

### Subtype-specific transcriptional dysregulation is dominated by downregulated genes and is most pronounced in AD4

To characterize the cell-type-resolved molecular basis of these subtypes, we compared differentially expressed genes (DEGs, relative to control) across MCI and AD1-AD4 in oligodendrocytes (ODC), microglia (MIC), and astrocytes (ASC) (Fig. 7d). Complementary one-vs-rest Wilcoxon analysis identified the top subtype-enriched genes for each AD subtype (Supplementary Fig. 7a). Across all three cell types and nearly all subtypes, downregulated DEGs vastly outnumbered upregulated DEGs (e.g., ODC: 1,293 down vs. 124 up in AD4; ASC: 775 down vs. 216 up in AD4; MIC: 1,036 down vs. 55 up in AD4), indicating that transcriptional suppression is the dominant signature of subtype-associated molecular pathology. Sequencing depth was about two-fold higher in AD2 and AD4 than AD1 and AD3, which may inflate DEG counts in deeper-sequenced subtypes. This alone, however, does not explain the AD4-predominant downregulation, as AD2’s comparable depth did not yield a similarly large DEG burden. AD4 carried the largest overall DEG burden in ODC and ASC, with the majority of these genes subtype-specific rather than shared with AD1-AD3 or MCI. Microglia were a notable exception to the down-dominant pattern: AD3 showed a large, largely subtype-specific upregulated signature (216 genes, one of the largest “Up” set of any subtype/cell-type combination) alongside a comparatively modest downregulated set, distinguishing AD3’s microglial response from the strongly down-dominant profiles of AD2 and AD4 (998 and 1,036 downregulated genes, respectively). Together, these data indicate that each AD subtype is associated with a largely distinct, cell-type-specific transcriptional program, with limited overlap in affected genes across subtypes.

### Cell-type-specific co-expression module activity defines four molecular AD subtypes

Because pooling all AD patients into a single group can mask whether a disease-associated signature is uniformly expressed or concentrated in a subset of individuals, we revisited the modules most strongly linked to AD status and to neuropathological/cognitive traits in each cell type (Figs. 2-3) and asked whether their eigengene expression differed systematically across the four molecular subtypes. To characterize molecular heterogeneity within AD, we examined eigengene expression of cell-type-specific co-expression modules (ASC-M5, ASC-M10, MIC-M5, MIC-M4, ODC-M1, EXC-M1, EXC-M2, INH-M7, INH-M10) across Control, MCI, and the four AD subtypes (Fig. 8a; Supplementary Fig. 7b). The subtypes showed clearly distinguishable module signatures, though they are not a separate analysis but a recombination of the shared architecture. AD2 and AD4 were characterized by elevated astrocyte and microglial module eigengenes (ASC-M5, ASC-M10, MIC-M4) relative to Control, consistent with heightened glial activation in these subtypes. Both ASC-M5 and ASC-M10 were selectively elevated in AD2 and reached their highest expression in AD4, consistent with activation of astrocyte remodeling and growth factor signaling programs in the Tissue Remodeling subtype, and ODC-M1 expression declined across AD3 and reached its lowest levels in AD4, consistent with disruption of oligodendrocyte lipid homeostasis in this subtype. AD3, in contrast, showed a marked depression of excitatory neuron modules (EXC-M1, EXC-M2), with eigengene values falling well below Control and all other AD subtypes, indicating a distinct pattern of neuronal dysfunction. AD4 additionally showed elevated inhibitory neuron module eigengenes (INH-M7, INH-M10) alongside its glial and excitatory neuron signatures, suggesting a broadly hyperactive or compensatory transcriptional state. Although INH-M7 was overall downregulated in AD relative to controls, its highest eigengene levels were observed in AD4, highlighting heterogeneity in interneuron-associated stress and inflammatory programs across AD. AD1 module eigengenes most closely resembled Control and MCI across nearly all modules, suggesting it represents a milder or earlier-stage molecular profile relative to AD2-AD4. The metal-ion homeostasis programs captured by INH-M1 (upregulated in AD) and EXC-M6 (downregulated in AD, suggesting loss of a protective metal-handling and anti-aggregation program that may normally help maintain neuronal proteostasis; Fig. 2c, d) parallel the broader Metal-Ion Stress signature defining this subtype, and the pathology-independent microglial activation program MIC-M2 (Fig. 3b) conceptually parallels the pathology-cognition dissociation observed in AD1 (Fig. 7c).

### Subtype-specific reorganization of intercellular signaling pathways

To assess whether these module-level differences were accompanied by altered intercellular communication, we examined predicted cell-cell signaling for three pathways, ANGPTL, retinoic acid (RA), and TGFβ, across Control and each AD subtype (Fig. 8b, Supplementary Fig. 6a). In Control, AD1, and AD4, ANGPTL signaling emanated broadly from astrocytes to nearly all excitatory, inhibitory, and glial populations, and RA and TGFβ signaling were similarly intact, with astrocyte- and microglia-derived signaling reaching inhibitory (VIP, LAMP5, SST, PVALB) and excitatory (IT, L4 IT, L5/6 NP, L5/6 IT C3) populations. AD2 showed selective attenuation of ANGPTL signaling (reduced to a single ASC-L5/6 IT C3 connection) while RA and TGFβ signaling were largely preserved. AD3 showed near-complete loss of all three signaling pathways, with ANGPTL, RA, and TGFβ chord diagrams reduced to minimal or absent connections. This collapse of intercellular signaling in AD3 parallels its depressed excitatory neuron and oligodendrocyte module eigengenes (Fig. 8a), together suggesting that AD3 represents a subtype defined by broad breakdown of glia-to-neuron communication rather than glial activation per se, distinguishing it from the glial-activation-predominant AD2 and AD4 subtypes.

### CLUE/CMap analysis nominates largely non-overlapping candidate therapeutics across subtypes

To explore subtype-specific therapeutic vulnerabilities, we compared each subtype’s transcriptional signature against the CLUE/CMap perturbagen database. This generated a disease-state score for each compound (−1, signature-reversing/“treating“; +1, signature-mimicking/“promoting”) (Fig. 8c, Supplementary Table 4). Although CLUE/CMap is derived primarily from perturbation profiles generated in immortalized cancer cell lines rather than brain cells, it provides a systematic framework for prioritizing candidate therapeutics for subsequent experimental validation. Subtype-specific analyses revealed distinct perturbagen response profiles across the four AD subtypes. AD1 exhibited the most heterogeneous pattern of associations, with both signature-reversing compounds (e.g., OMDM-2, scopoline, perillyl-alcohol, and baricitinib) and signature-mimicking compounds (e.g., docetaxel, amineptine, and the HDAC inhibitors JNJ-26481585 and resminostat). However, none of AD1’s associated compounds met the significance threshold applied to the other subtypes, and these compounds are shown for comparison, not as significant nominations (Supplementary Table 4). AD2 was characterized by a smaller set of predominantly signature-reversing compounds, including rucaparib, pyroxamide, and semaxanib. AD3 yielded relatively few significant perturbagens overall, with signature-reversing compounds rosuvastatin, sappanone-a, and tetrabenazine. In contrast, AD4 displayed a broad signature-reversing profile across numerous compounds, including rucaparib, baricitinib, hydrocortisone, resminostat, JNJ-26481585, docetaxel, and lomustine. The limited overlap in perturbagen signatures across subtypes suggests that candidate therapeutic strategies may differ substantially among transcriptionally defined AD subtypes, and pharmacological strategies effective for one AD molecular subtype may not generalize to others, reinforcing the rationale for subtype-stratified therapeutic approaches.

### Stellar Atlas: an AI-native interface for community interrogation of the integrated atlas

A resource of this scale is only as valuable as it is accessible. Atlas-scale snRNA-seq studies are typically distributed as static supplementary tables or as gene-by-cell-type browsers that require a user to know in advance which gene, cell type, or contrast to inspect, leaving most of the integrated signal unexplored, especially for the experimental and clinical researchers best positioned to act on it. To remove this barrier and make the shared molecular architecture described above directly usable, we developed Stellar Atlas, an AI-native, conversational interface to the integrated atlas that is openly available both as a public web instance and as an installable software package (Fig. 9).

Stellar Atlas couples a Claude-backed conversational agent to the atlas’s live data stores, which include cell-type-resolved expression, differential expression results, co-expression modules with their hub-gene networks, and intercellular communication tables. Rather than selecting from a fixed gallery of precomputed views, a user poses questions in plain language, for example ‘which microglial modules track with Braak stage?’, ‘show the hub genes of the microglial lysosomal module’, or ‘how does EGF signaling change from control to AD?’, and the agent returns the relevant statistics, gene lists and figures computed on demand from the harmonized data, with an optional PubMed lookup for the surrounding literature. The gene regulatory networks and molecular subtypes reported here can be surfaced through the same module system as the resource develops.

Because the agent operates over the same harmonized data that underlies the analyses reported here, many of the figure-level results in this study can be reproduced and extended through the interface, and the agent’s answers are grounded in, and traceable to, the underlying tables rather than free text (Fig. 9). In this way Stellar Atlas turns a static atlas into a queryable, hypothesis-generating resource, allowing any investigator to interrogate the cell-type-specific programs and communication changes relevant to their own gene or pathway of interest without writing code.

## Discussion

In this study, we present an integrated single-nucleus atlas of the human cortex encompassing over 3 million nuclei from 791 individuals across 13 snRNA-seq datasets, representing a substantial advance in scale and analytical depth for AD single-cell research. Critically, our central advance is not scale per se but integration: by harmonizing the datasets we show that AD has a reproducible, shared molecular architecture, the same cell-type-specific modules, hub-TF regulatory networks and communication changes recur across datasets rather than appearing as study-specific signals. By applying a comprehensive multi-layer framework spanning co-expression network analysis, cell-cell communication inference, gene regulatory network reconstruction, sex-stratified transcriptomics, and deep generative modeling, we provide a systems-level view of AD pathobiology at unprecedented cellular resolution. Our findings establish that AD is not a uniform transcriptional program but a spectrum of cell-type-specific, sex-modulated, and individually heterogeneous molecular states, with important implications for disease understanding and therapeutic development.

The co-expression network analyses revealed that each major cortical cell type harbors distinct co-expression modules with specific associations with disease severity metrics. Among non-neuronal cells, the oligodendrocyte module M4 exhibited the strongest AD-associated transcriptional shift and was enriched for genes involved in myelin maintenance and neuronal support, despite not being significantly correlated with clinical and neuropathological traits within the ROSMAP cohort. These findings are consistent with emerging evidence that white matter and myelin disruption are prominent features of AD and may contribute to disease pathogenesis even when such changes are not reflected by conventional neuropathological staging measures. Similarly, the identification of lysosomal, protein-clearance, and immune-signaling modules in microglia that correlate with disease severity underscores the centrality of microglial dysfunction in the management of amyloid and tau pathology. The observed female-biased microglial immune-activation program is consistent with prior reports of sex-dependent immune responses in AD, including effects linked to X chromosome dosage, escape from X-chromosome inactivation, and *APOE*-dependent modulation of microglial activation.

The intercellular analysis provides a new dimension to our understanding of AD progression by revealing how intercellular communication networks are systematically remodeled across disease stages. The finding suggests that MCI represents not merely a transitional state but a phase with molecularly distinct intercellular signaling that may be targetable prior to the onset of frank neurodegeneration. The loss of EGF signaling in AD aligns with established findings of loss of growth-factor-mediated communication between neuronal sub-cell types and other neuronal and glial cell types in AD.

Our sex-stratified analyses provide some of the most comprehensive evidence to date that AD is a sexually dimorphic disease at the transcriptional regulatory level. The identification of sex-biased regulons and TF-TG regulatory edges across multiple cell types suggests that female and male brains respond to AD pathology through overlapping but distinct molecular programs. This has direct implications for clinical trial design and drug development: therapies optimized in predominantly male or sex-unbalanced cohorts may have differential efficacy in female patients. Integrating sex as a biological variable in MONET, and adjusting for sex as a covariate in pseudobulk profiles prior to subtyping, ensures that the four AD subtypes we identify reflect molecular heterogeneity that is not confounded by sex effects.

The four AD subtypes identified by MONET represent a robust and biologically interpretable molecular classification of AD individuals. The association of each subtype with distinct pathway signatures (metal ion stress, neuroinflammation, synaptic integrity, and tissue remodeling) suggests that different AD patients may be primarily driven by distinct pathological mechanisms, a hypothesis with important implications for precision medicine. Drug repurposing analysis through the CLUE database nominated specific therapeutic classes for each subtype, with HDAC inhibitors showing cross-subtype relevance, suggesting both subtype-specific and shared therapeutic opportunities. The module-trait correlations in this study (Figs. 2 and 3) were established within the ROSMAP cohort, which provided complete and uniformly coded clinical annotation; whether these gene module-pathology relationships replicate in independent cohorts such as SEA-AD remains to be tested. This contrasts with the subtype clinical characterization in Fig. 7c, which drew on both cohorts because the subtypes themselves were defined from pooled multi-cohort data; there, CERAD scores were harmonized across cohorts’ differing scoring conventions prior to comparison.

One of the most notable findings of this study is the apparent dissociation between classical neuropathological burden and cognitive impairment across transcriptionally defined AD subtypes. In particular, the Metal Ion Stress subtype (AD1) exhibited disproportionately severe cognitive impairment relative to its neuropathological burden. Although this observation will require validation in independent cohorts and adjustment for potential confounders (including age, *APOE* genotype, post-mortem interval, cohort composition, coexisting neuropathologies, and sample size), it raises the possibility that molecular mechanisms not fully captured by conventional amyloid and tau pathology contribute to clinical severity in this subtype.

Limitations include reliance on post-mortem cross-sectional data, the lack of longitudinal information to track subtype stability, and limited representation of endothelial and oligodendrocyte precursor cells. A further limitation is that the four molecular subtypes, although reproducible across random seeds, were derived and evaluated within the same integrated cohort collection. Independent external validation, ideally in prospectively collected cohorts, will be required to establish their stability, generalizability, and clinical utility while excluding residual cohort-specific effects. The atlas and MONET framework are designed to be projectable: because subtyping operates on a covariate-adjusted, cell-type-resolved feature space rather than a dataset-specific embedding, new donors can in principle be mapped onto the existing latent space and evaluated against the identified subtypes. Large independent resources such as the PsychAD consortium, which aggregates multiple cohorts including MSSM, HBCC, and RADC, provide an opportunity to test the generalizability of these transcriptional subtypes, offering a direct route to the external validation called for above^28–34^. Future work integrating spatial transcriptomics, longitudinal cohorts, and functional validation of key regulatory nodes will be essential to translate these findings into clinical utility. Nevertheless, this atlas establishes a foundational resource and analytical framework for the field, advancing our understanding of AD heterogeneity and providing actionable hypotheses for mechanistic and therapeutic investigation. Finally, by releasing the integrated atlas together with Stellar Atlas, an AI-native conversational interface, we aim to make this shared molecular architecture not only a static reference but a living, queryable resource that the community can interrogate directly, lowering the barrier between a large multi-cohort dataset and the specific, testable hypotheses that individual laboratories need.

## Methods

### Datasets and data access

Thirteen publicly available snRNA-seq datasets derived from human prefrontal cortex and temporal cortex were included in this study. Datasets span three diagnostic categories: Alzheimer’s disease (AD), mild cognitive impairment (MCI), and cognitively normal controls. Diagnosis labels were harmonized across the heterogeneous nomenclature used by the 13 contributing cohorts by retaining each cohort’s own diagnostic classification: donors with a cohort-reported diagnosis of cognitively normal/no dementia were labeled Control, donors with mild cognitive impairment were labeled MCI, and donors with any other cohort-reported dementia-spectrum diagnosis (e.g., AD, AD with Down syndrome, AD with vascular dementia, vascular dementia) were labeled AD. After integration, the atlas comprised data from 791 individuals. All data were accessed from public repositories (GEO, Synapse) under their respective data use agreements. Dataset-level metadata, including tissue regions, sequencing platforms, and sample sizes, are provided in Supplementary Table 1.

### Quality control and doublet removal

Raw count matrices for each dataset were processed independently. Each dataset underwent an initial global QC filtering step, removing ambient RNA (CellBender v0.3.0), retaining cells with at least 200 genes per nucleus and a total count of at least 250 transcripts^35^. Quality control was performed using Scanpy^36^ . Genes detected in at least 3 cells/nuclei were retained, and no more than 15% mitochondrial gene content was allowed. Prior to dataset merging, doublets were identified and removed using Scrublet with default parameters and dataset-specific expected doublet rates^37^. After QC, datasets were merged and subjected to integration (Supplementary Fig. 1).

### Data integration and cell type annotation

Datasets were integrated using scVI, with study identity as the batch covariate ^38^. Reference-based cell type annotation was performed using scANVI with the Allen Brain Atlas single-cell reference ^39, 40.^ Leiden clustering was applied to the integrated latent space, yielding 17 clusters. Clusters were annotated to seven major cell type classes (EXC, INH, ODC, OPC, ASC, MIC, ENDO) based on expression of canonical marker genes and scANVI label predictions^39^. UMAP dimensionality reduction was computed using the scVI latent representation for visualization^38^.

### Differential expression analysis

Differential expression (DE) analysis for each celltype in AD, MCI, Control was performed using the MAST framework^41^ (Model-based Analysis of Single-cell Transcriptomics) implemented via Seurat’s FindMarkers function (test.use = “MAST”). Prior to testing, raw counts were log-normalized using a library-size scale factor of 10,000 (NormalizeData, normalization.method = “LogNormalize”). All genes were tested with no minimum expression filter (min.pct = 0) and no log-fold-change pre-filter (logfc.threshold = 0), retaining the full gene list for downstream significance filtering. Disease-stage comparison: To identify cell-type-resolved transcriptomic signatures of disease, single-cell MAST DE was performed for three pairwise disease-stage comparisons - AD vs. Control, MCI vs. Control, and AD vs. MCI - separately for each of the cell types: astrocytes, excitatory neurons, inhibitory neurons, microglia, oligodendrocytes. Seurat objects containing the full complement of cells per major type (HVG-restricted RNA assay) were used as input. MCI cells were excluded from the AD vs. Control comparison. Significant DEGs were defined as adjusted p-value < 0.05 (Bonferroni correction applied by Seurat over all tested genes). Sex-stratified comparison: Sex-stratified DE analysis was performed to identify sex-biased transcriptomic programs in the context of AD. Within each disease stage (Control, MCI, Dementia) and each major cell type, cells were subset to that stage and MAST DE was performed comparing male versus female donors (ident.1 = Male, ident.2 = Female). Positive avg_log2FC values indicate higher expression in males. AD-subtype comparison: To characterize transcriptomic differences associated with each AD molecular subtype, MAST DE was performed comparing each subtype (AD1, AD2, AD3, AD4) and MCI independently against the Control group, within each major cell type. Sample-level subtype assignments (AD1-AD4, MCI, Control) derived from GMM clustering of the VAE latent space (see AD subtyping section) were propagated to individual cells via their donor sample identifier; cells from donors without a valid subtype label were excluded from analysis. For each comparison, cells from only the two groups under test were retained prior to normalization and testing. Positive avg_log2FC values indicate higher expression in the test group (subtype or MCI) relative to Control. In total, 35 DE tests were performed (5 comparisons × 7 cell types). Significant DEGs were defined as adjusted p-value < 0.05. Overlap of DEG signatures across subtypes within each cell type was visualized using UpSet plots (upsetplot Python package), with a reporting threshold of adjusted p-value < 0.05 and |log2 fold change| > 1; intersection sets containing fewer than 5 genes were suppressed.

### Hierarchical weighted gene co-expression network analysis

Weighted gene co-expression network analysis adapted for single-cell data (hdWGCNA) was performed separately for each major cell type. Metacells were constructed by aggregating a minimum of 25 neighboring cells in the shared nearest-neighbor graph. Soft-thresholding powers were selected using the scale-free topology criterion (R-squared > 0.80). Co-expression modules were identified using hierarchical clustering of the topological overlap matrix (TOM). Module eigengenes (MEs) were computed as the first principal component of within-module gene expression variation across metacells. Module-trait correlations were computed using Spearman rank correlation between per-sample module eigengenes and clinical variables (Braak stage, CERAD score, MMSE), with FDR correction for multiple comparisons. Differential module eigengene (DME) analysis compared ME scores between AD and control using linear mixed models with donor as a random effect, adjusted for age, sex, and study. Only module-trait correlations passing the multiple-testing threshold (FDR < 0.05) are displayed. Braak stage, CERAD score, and MMSE values used in these correlations were obtained from the ROSMAP cohort (n=176), which provided complete and uniformly coded pathological and cognitive annotation across all samples with co-expression modules.

### Cell-cell communication inference

Intercellular communication was inferred using CellChat v2^42^, applied to log-normalized single-cell expression matrices, with separate objects built for each state (Control, MCI, AD) and differential interaction analysis was carried out for pairwise comparisons: MCI vs. Control, AD vs. MCI, and AD vs. Control. Signaling pathways were ranked by differential interaction strength, and pathways with significant changes (permutation test p < 0.05) were retained for visualization and interpretation.

### Gene regulatory network inference (pySCENIC and NetREm)

Gene regulatory network inference was performed using pySCENIC^24^. For each cell type, the pySCENIC pipeline was executed in three stages: (1) co-expression module identification using GRNBoost2 on the single-cell gene expression data, (2) TF motif enrichment using cistarget databases to prune modules to high-confidence regulons, and (3) regulon activity scoring per cell using AUCell. Regulon activity scores were averaged per sample and correlated with disease trait variables. To prioritize high-confidence TF-target gene (TG) interactions, pySCENIC regulons (which formed our “prior GRN”) were further refined using NetREm, a network-constrained regression approach that leverages protein-protein interaction (PPI) network topology to regularize TF-TG interaction weights. We applied NetREm for a given cell type using the following inputs [single-cell gene expression data, PPI network (“Main NetREm PPIN” used in the NetREm study that was derived from several PPI databases of direct and/or indirect interactions including STRING database version 12) as prior knowledge of TF-TF interactions, prior GRN (predicted by pySCENIC)] and parameters [network-constrained prior term *β* = 100 and sparsity prior term *α* selected from Scikit-Learn’s LassoCV model with 5-fold cross-validation (CV)] ^43^.

The combined pySCENIC and NetREm framework yielded cell-type-specific GRNs with high-confidence directional TF-TG edges. Regulon activity was scored per cell using AUCell, and regulon activity scores were correlated with disease traits across samples. NetREm complemented pySCENIC by applying network-constrained regularization to prioritize TF-TG interactions supported by both co-expression and PPI network topology, and additionally resolved TF-TF coordination for TG co-regulation as either cooperative (TFs jointly engaging shared targets) or antagonistic (TFs with opposing regulatory effects on shared targets); thus, NetREm output two types of TF-TF coordination networks for a given cell-type: TG-specific and overall. Sex-biased regulons were identified by comparing AUCell-scored regulon activity between female and male AD donors within each cell type.

### Pseudobulk generation and covariate-adjusted expression profiling

Single-cell RNA-sequencing data were aggregated into sample-level pseudobulk expression profiles in a cell-type-specific manner. For each cell type, raw count matrices were extracted from Seurat objects, with genes as rows and individual cells as columns. Cell-level metadata were used to assign cells to biological samples, and gene-level counts were summed across all cells belonging to the same sample to generate gene-by-sample pseudobulk expression matrices independently for each cell type. Pseudobulk expression matrices were aligned to sample-level metadata and restricted to samples present in both expression and covariate datasets. Expression values were log2-transformed following addition of a pseudocount. To account for technical and demographic confounders, linear regression was performed independently for each gene to remove effects of study, age, and sex. Regression coefficients were estimated using bootstrap resampling (1,000 iterations), and the median coefficient across bootstrap replicates was used for adjustment. Covariate-adjusted expression values were obtained by subtracting estimated contributions of study, age, and sex from the original expression values. PCA and correlation analyses were performed before and after regression to verify effective confounder removal.

### Construction of cell-type-specific gene expression vectors

Covariate-adjusted pseudobulk expression matrices from seven cell types (astrocytes, microglia, endothelial cells, oligodendrocytes, OPCs, inhibitory neurons, and excitatory neurons) were integrated at the gene-sample level. For each gene and sample, cell-type-specific expression values were aggregated into a fixed-length vector of seven elements in a predefined and consistent cell-type ordering. Only vectors containing exactly seven positions were retained. Missing values arising from absent cell-type measurements were explicitly encoded.

### Missing value imputation, masking, and vector padding

A position-specific imputation strategy was applied. For each gene and each position within the seven-element vector, the mean expression value across all samples with observed data was computed. Vectors with no missing values were retained unchanged; vectors with exactly one missing value were retained after replacing the missing entry with the corresponding gene- and position-specific mean. The resulting imputed value was thereafter treated identically to a directly observed value. Genes with more than one missing value were excluded. Gene-specific vectors were concatenated across all genes to form sample-level feature vectors, then zero-padded to a fixed length of 35,000 features; a binary mask was generated to distinguish real gene positions (1, whether directly observed or mean-imputed) from the zero-padded tail (0), yielding uniformly sized sample-level representations.

### Masked variational autoencoder architecture

A VAE was implemented within the MONET framework to learn a low-dimensional latent representation of each sample. The encoder comprised six fully connected layers (input, 8,192, 4,096, 2,048, 1,024, 512, and 256 units) with ReLU activations and dropout (p = 0.30) after the first two layers. Two independent linear projections from the 256-dimensional encoder output produced the mean (*μ*) and log-variance (log *σ*^2^) of a 128-dimensional Gaussian latent distribution. Latent vectors were sampled via the reparameterization trick, *z* = *μ* + *σε*, where *ε*∼ N(0, I). The decoder mirrored the encoder (128, 256, 512, 1,024, 2,048, 4,096, 8,192, and input dimension), with ReLU activations on all layers except the final output layer.

### Training procedure

The VAE was trained independently across 20 random seeds (seeds 0-19). For each seed, samples were split 80/20 into training and held-out sets; all samples were subsequently embedded using the trained encoder and used for downstream clustering. The model was optimized using Adam (learning rate =1 × 10^−4^, batch size = 32) for up to 200 epochs. The training objective combined masked reconstruction loss with Kullback-Leibler (KL) divergence:

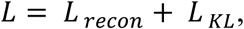

Where the masked reconstruction loss was

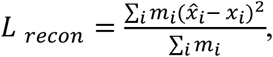

And the KL-divergence term was

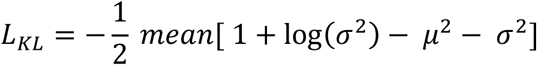

where *m_i_* is the binary mask excluding only the zero-padded tail of each sample vector from the reconstruction objective; mean-imputed positions were retained and contributed gradient on the same basis as directly observed values. Gradient norms were clipped to a maximum of 1.0. The model checkpoint with the lowest training loss (improvement threshold delta = 1 × 10^−4^) was retained.

### GMM clustering and ensemble consensus

After training, latent representations were obtained for each sample *i* using the best-checkpoint encoder: the encoder produced the mean and log-variance (*μ_i_*, log *σ*^2^), and a latent embedding *z_i_* = *μ_i_* + *σ_i_* × *ε_i_* was drawn via the reparameterization trick. For each of the 20 seeds, a GMM with *k* = 4 was fitted to these latent embeddings *z_i_*. Silhouette analysis was performed across candidate values of *k* (*k* = 2-10) and favored a two- or three-cluster solution; nevertheless, a four-cluster solution was selected and applied uniformly across all seeds, to balance cluster separation with increased resolution of molecular heterogeneity. A symmetric *N* × *N* pairwise co-assignment matrix *M* was constructed as

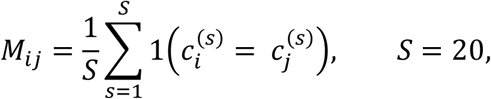

where 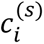 denotes the GMM cluster assignment of sample *i* for seed *s*, and **1**(⋅) is the indicator function. Thus, *M_ij_* represents the fraction of seeds in which samples *i* and *j* were assigned to the same GMM cluster. The consensus distance matrix was defined as

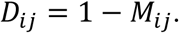

Agglomerative hierarchical clustering with average linkage was applied to *D*, and the resulting dendrogram was cut at *k* = 4 to obtain the final consensus cluster assignments (Supplementary Fig. 4).

### Seed-to-consensus alignment and stability assessment

Each seed’s cluster labels were aligned to the consensus using the Hungarian algorithm (scipy.optimize.linear_sum_assignment), which maximizes overlap between seed-specific and consensus assignments. Per-sample agreement fractions were computed as the proportion of seeds (after alignment) in which the sample’s cluster matched its consensus assignment. Clustering stability was further assessed by computing within-cluster and between-cluster co-assignment distributions. The block-diagonal structure of M ordered by final cluster assignment confirmed that within-cluster co-assignment substantially exceeded between-cluster co-assignment. Each SEA-AD sequencing library was treated as an independent input sample for pseudobulk feature vector construction and subtyping; donors with multiple independently sequenced libraries or represented in more than one constituent dataset (ROSMAP, Mathys et al. 2019, and Zhou et al. 2020) were retained as separate entries under their library or dataset of origin. Among the 391 samples assigned a subtype, 21 (5.4%) belong to donors with more than one sample in the input whose duplicate samples received different subtype calls; 13 of 36 SEA-AD donors with replicate libraries (36.1%) and 8 of 12 comparable cross-dataset donors (66.7%). This reflects sample-level, not donor-level, subtyping: the overall four-subtype structure remained stable across 20 independent seeds (Fig. 7b), and donor-level assignment for this subset of samples is an area of continued refinement.

### Sankey visualization

Sankey diagrams were generated using Plotly to visualize the mapping from each seed’s GMM cluster assignments to the final consensus clusters. Individual per-seed diagrams and a combined 4×5 grid view across all 20 seeds were produced. Link widths are proportional to the number of samples flowing from each seed cluster to each consensus cluster (Supplementary Fig. 5).

### Clinical trait comparison across subtypes

Braak stage, CERAD score, and MMSE were compared across the four consensus AD subtypes using the subset of ROSMAP^10^ and SEA-AD^14^ samples with complete clinical annotation (n=217; ROSMAP n=176, SEA-AD n=41). Because the four subtypes were derived from pooled multi-cohort pseudobulk data and are not evenly distributed across cohorts, both cohorts were retained for this comparison. ROSMAP and SEA-AD score CERAD neuritic plaque density using opposite numeric conventions: ROSMAP’s ceradsc variable is coded 1 (Definite AD, most severe) to 4 (No AD, least severe), whereas SEA-AD’s CERAD score follows the standard convention of 0 (Absent, least severe) to 3 (Frequent, most severe). To enable direct comparison across cohorts, ROSMAP CERAD scores were standardized onto SEA-AD’s scale by inversion (CERAD_standardized = 4 − CERAD_raw); SEA-AD scores were retained unchanged. Braak stage required no such correction, as both cohorts use the same direction and range (0-6). Pairwise differences in Braak stage, harmonized CERAD score, and MMSE were assessed using two-sided Mann-Whitney U tests with Benjamini-Hochberg correction for multiple testing (Fig. 7c; Supplementary Fig. 6b).

### Drug repurposing analysis (CLUE/CMap)

Drug repurposing candidates for each AD subtype were identified using the Connectivity Map (CLUE) database. Subtype-specific differentially expressed gene signatures (AD vs. control, within each subtype) were generated using the pseudobulk MAST framework. The top differentially expressed genes (ranked by signed − log_10_(FDR)) were submitted as query signatures to the CLUE API. Drug-subtype connectivity scores were computed, and results were filtered by FDR and absolute connectivity score thresholds. Drug classes nominated for each subtype are reported together with the highest observed significance (Subtype 2: − log_10_(FDR) = 15.65).

### Stellar Atlas interface

Stellar Atlas is the public deployment of STELLAR (Single-cell Transcriptomic Exploration with LanceDB and Atlas Retrieval), an open-source framework we developed to turn a processed AnnData (h5ad) or Seurat object into a deployable, interactive snRNA-seq web atlas (https://github.com/swaruplab/stellar-atlas, MIT license). On ingestion the processed atlas is written to two complementary stores. A gene-major LanceDB store holds the cell-by-gene matrix with one row per gene and sparse cell-index and value columns, so that recoloring a UMAP by any gene is a single-row read rather than a scan across millions of cells. A DuckDB store holds all non-expression tables, including cell and donor metadata, differential expression results, hdWGCNA modules, CellChat networks and pseudobulk precomputes, in one embedded columnar file that returns analytical aggregates over millions of rows in milliseconds.

A FastAPI service exposes the routes consumed by a single-page React application, and the interface is modular, so that only the analyses enabled for a project appear. For the panAD atlas these comprise UMAP and per-cell-type expression and violin views, the differential expression viewer with volcano plots and lasso-to-enrichment, the hdWGCNA co-expression modules with hub-gene networks and DME heatmaps, and the CellChat chord diagrams and ligand-receptor tables. Natural-language interaction is provided by a Claude-backed conversational agent (the copilot module) that is given tool access over the same live atlas stores, so that its answers are computed from the underlying data rather than recalled from text, together with an optional PubMed literature-lookup tool scoped to neurodegeneration. The deployed panAD instance is available at https://swaruplab.bio.uci.edu/panad_atlas/, and the full source, documentation and configuration schema are available at https://github.com/swaruplab/stellar-atlas under an MIT license.

### Software and statistical analysis

Analyses were performed using Python (version 3.10) and R (version 4.3). Key Python packages included PyTorch (VAE training), scikit-learn (GMM, hierarchical clustering, silhouette analysis), SciPy (Hungarian algorithm, statistical tests), Plotly (Sankey visualization), pySCENIC^23^(gene regulatory network inference), NetREm^25^ (network-constrained regression), and Scanpy^36^ (single-cell data processing and Wilcoxon rank-sum differential expression). R packages included Seurat^44^ (single-cell data handling), hdWGCNA^22^ (co-expression network analysis), MAST^41^ (differential expression), and CellChat (cell-cell communication). All statistical tests were two-sided unless otherwise noted. Multiple testing correction was performed using the Benjamini-Hochberg FDR method.

## Data availability

All snRNA-seq datasets used in this study are publicly available. Supplementary Table 1 provides the corresponding studies and PubMed identifiers and/or data repository accessions. The integrated atlas, processed data, metadata, differential expression results, and cell-type-resolved GRN data (NetREm- and pySCENIC-derived TF-target gene edges, Fig.5) generated in this study are available at https://swaruplab.bio.uci.edu/panad_atlas/. This integrated atlas can be interactively explored through Stellar Atlas, an AI-native, conversational interface to the panAD dataset, openly available as a public web instance https://swaruplab.bio.uci.edu/panad_atlas/ and an installable software package https://github.com/swaruplab/stellar-atlas.

## Code availability

All analysis code, including the MONET framework, pseudobulking workflows, hdWGCNA wrapper scripts, CellChat analysis scripts, pySCENIC, and NetREm pipelines, is available at https://github.com/neginrhm/panAD-MONET under the MIT license.

## Acknowledgements

We thank Luis Solano, Nellie Kwang, Kristen Vallejo, and members of the Swarup laboratory for their helpful comments on the manuscript. We also thank the Research Cyberinfrastructure Center (RCIC) at the University of California, Irvine for providing high-performance computing resources and technical support. This work was supported by the Alzheimer’s Association Research Grant (AARG-25-1472103), the Cure Alzheimer’s Fund, and the National Institute on Aging of the National Institutes of Health under award numbers R01AG071683 and U54AG054349 to V.S. N.R. was supported by the National Institutes of Health under award number 1T32GM136624-01.

## Author Contributions

N.R., S.M. and V.S. conceptualized this study. N.R. and V.S. wrote the manuscript with input from and approval by all authors. S.M. and Z.C. downloaded individual snRNA-seq datasets and assisted with data quality control and integration. N.R. generated the panAD atlas, performed the bioinformatics analyses, and developed the subtyping workflow. S.K. performed NetREm and associated GRN analyses. N.R. performed pySCENIC analysis and completed the GRN analysis. V.S. generated the panAD and Stellar Atlas websites. S.M., Z.S., S.K., and Z.C. provided guidance and edits on the manuscript.

