## Supplementary Figures for "An Integrated Single-Nucleus Atlas Resolves Cell-Type-Specific Programs and Molecular Subtypes in Alzheimer’s Disease"

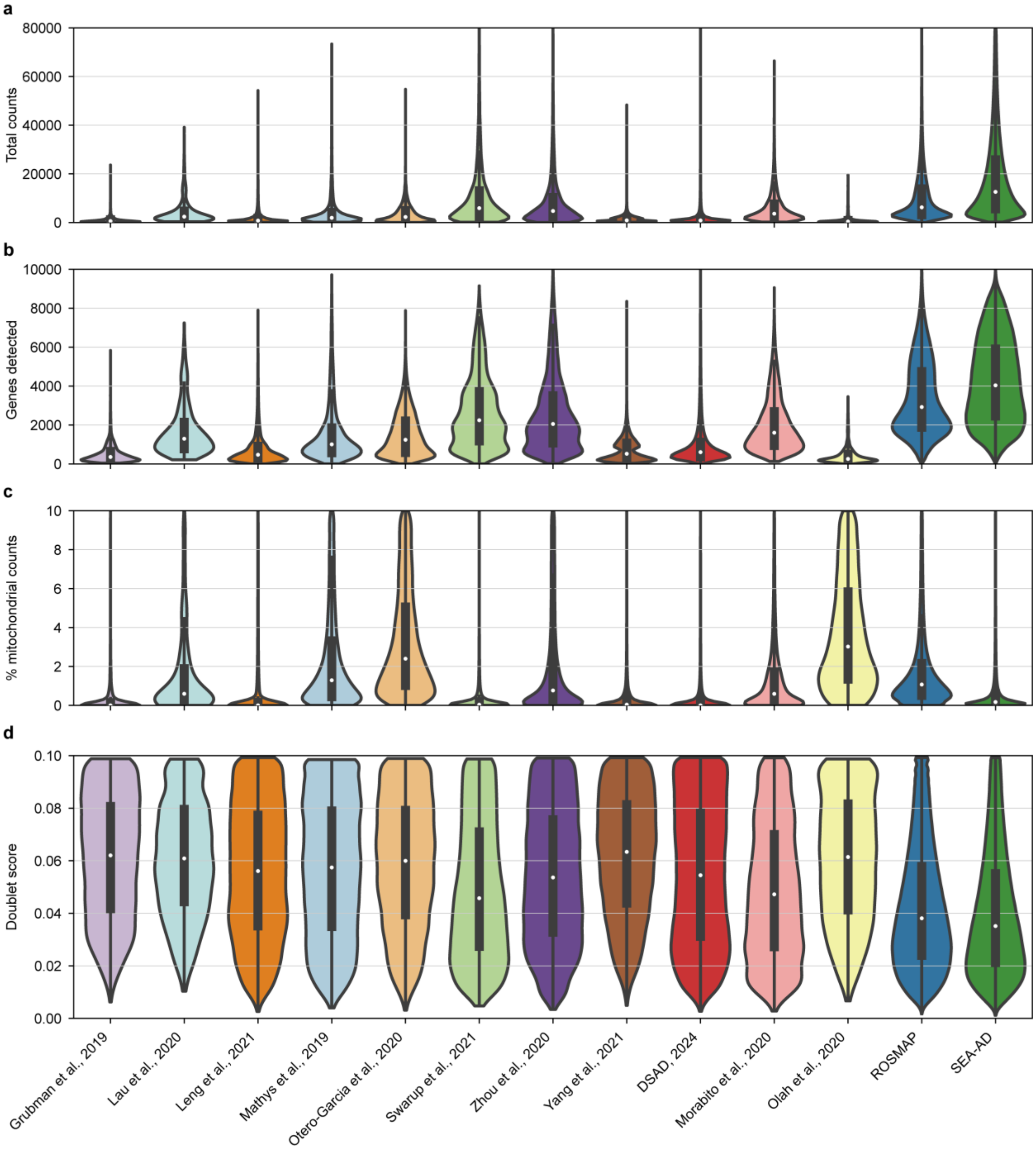


**Supplementary Figure 1 |** Per-dataset single-nucleus/single-cell RNA-sequencing quality-control metrics. **a-d,** Distribution across nuclei of **a,** total UMI counts, **b,** number of genes detected, **c,** percentage of mitochondrial transcript counts, and **d,** doublet score, shown separately for each of the 13 constituent studies.

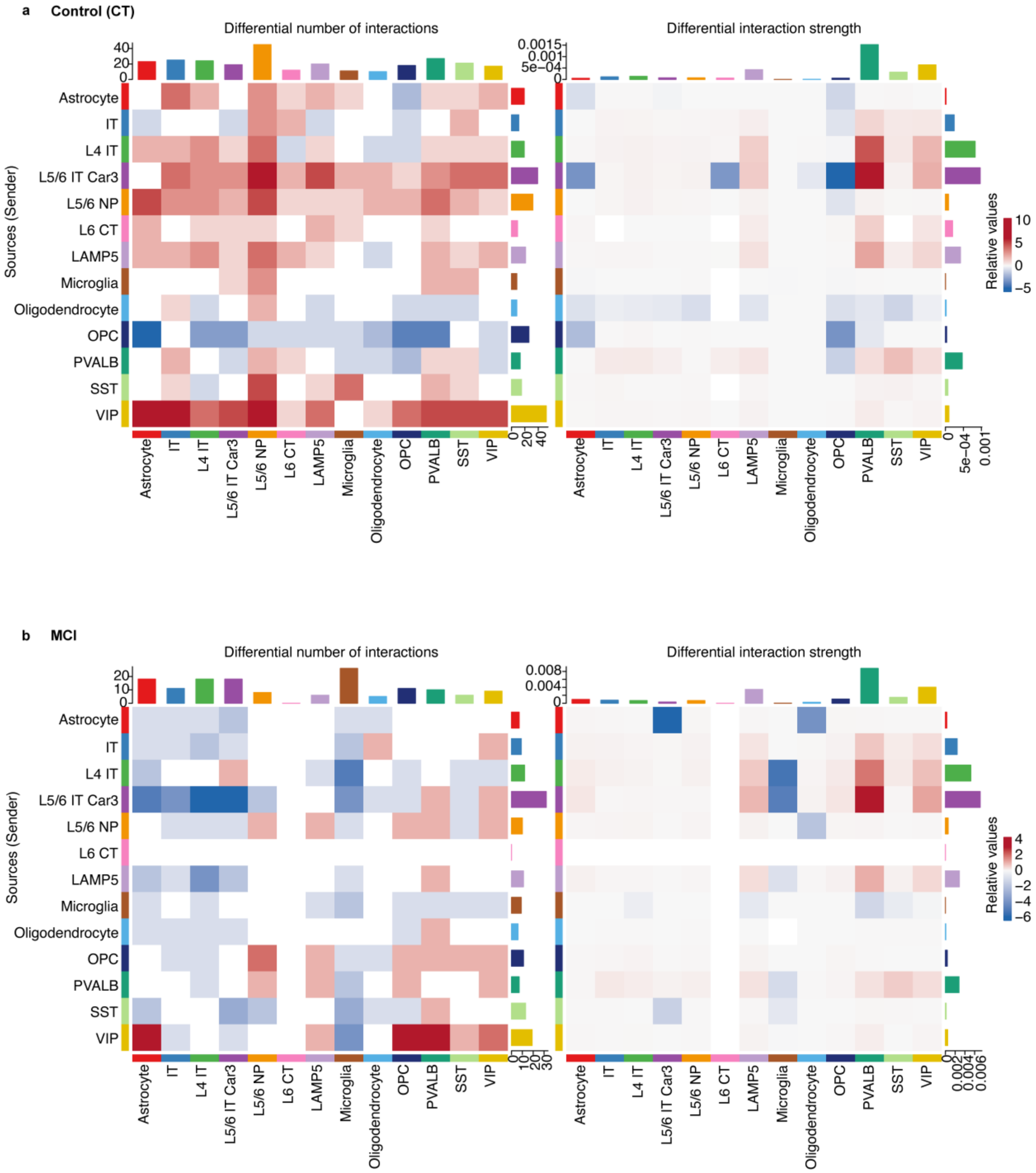


**Supplementary Figure 2 |** Sex-stratified cell-cell communication across diagnostic groups. **a,b,** CellChat differential number of interactions (left heatmap) and differential interaction strength (right heatmap) between female and male donors across cell types, shown separately for **a,** cognitively normal controls (CT) and **b,** mild cognitive impairment (MCI). Red indicates interactions relatively increased in male donors; blue indicates interactions relatively increased in female donors. Bar plots along the top and right margins of each heatmap show the total differential signaling per cell type.

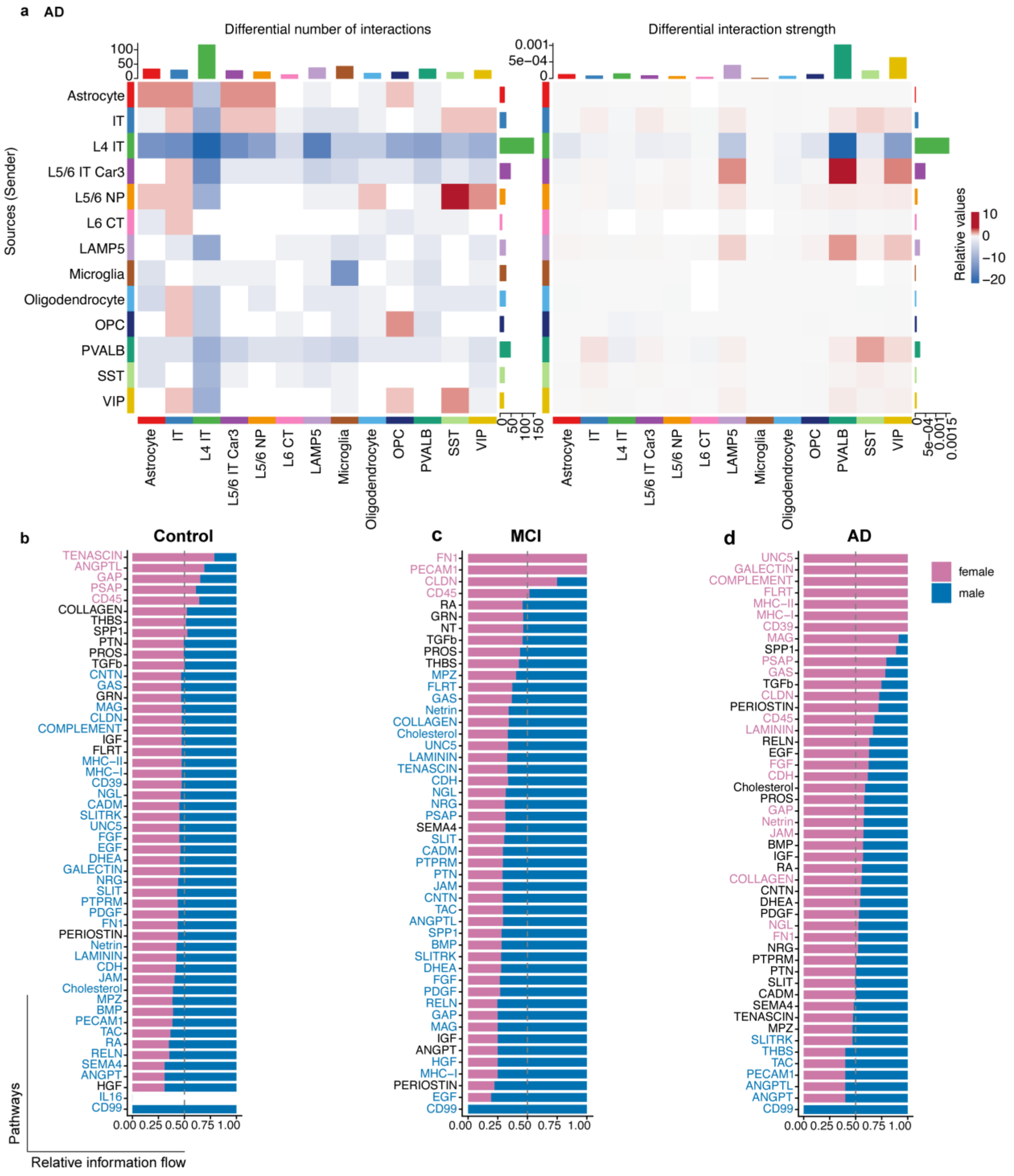


**Supplementary Figure 3 |** Sex-stratified cell-cell communication and signaling pathway information flow in AD. **a,** CellChat differential number of interactions (left) and differential interaction strength (right) between female and male donors across cell types in Alzheimer's disease (AD), plotted as in Supplementary Fig. 2. **b-d,** Relative information flow of each detected signaling pathway between female (pink bars) and male (blue bars) donors in **b,** cognitively normal controls (CT), **c,** MCI, and **d,** AD. Pathway name labels are colored pink or blue where information flow differs significantly between female and male donors (colored by the sex with relatively higher flow) and black where no significant sex difference was detected.

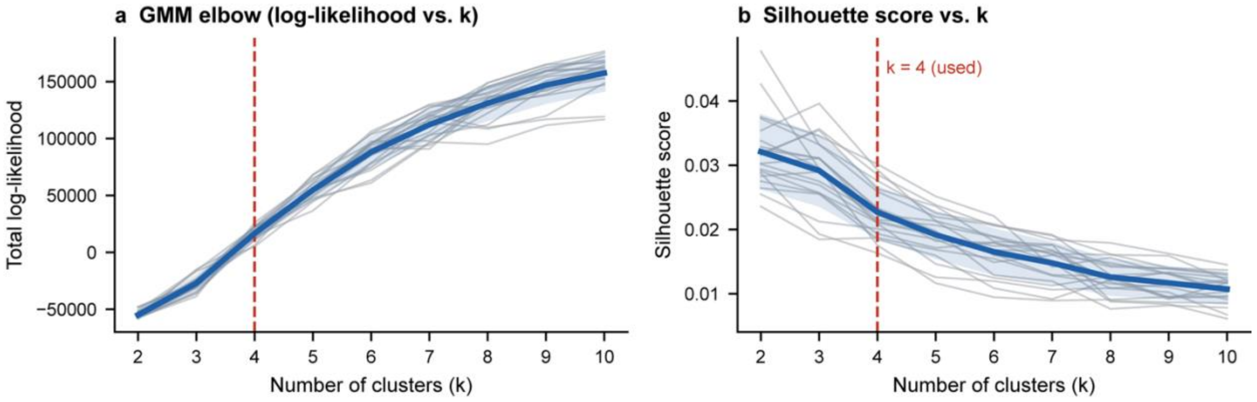


**Supplementary Figure 4 |** GMM cluster-number diagnostics across all 20 training seeds. **a,** Total Gaussian mixture model (GMM) log-likelihood as a function of the number of clusters (k), fit independently for each of 20 MONET training seeds. **b,** Silhouette score as a function of k. Thin gray lines, individual seeds (n = 20); blue line and shaded band, mean ± s.d. across seeds. Dashed red line marks k = 4, the value retained for downstream subtyping despite silhouette score favoring k = 2-3 (see main text for rationale).

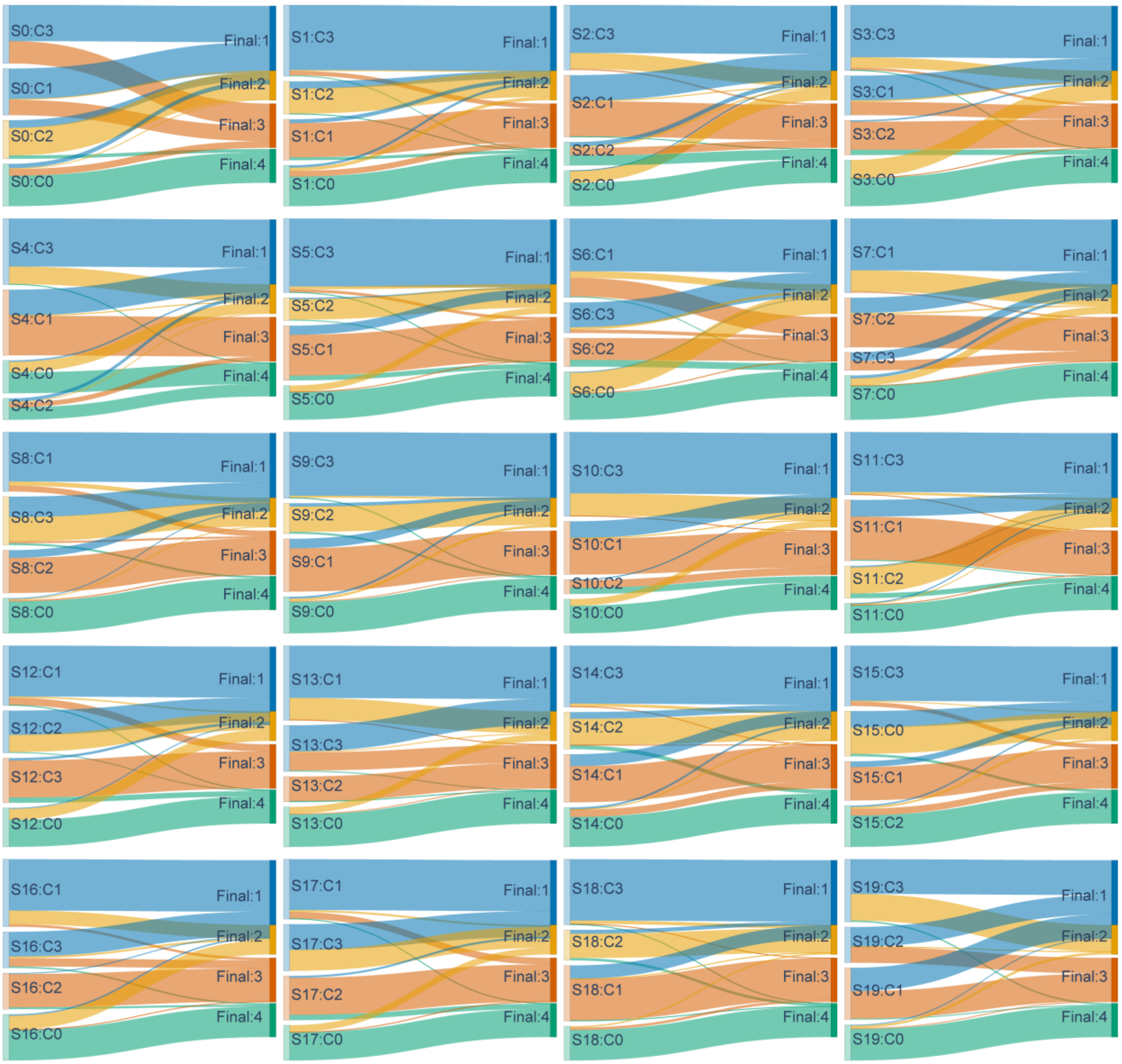


**Supplementary Figure 5 |** Consensus subtype assignment across all 20 clustering seeds. Sankey diagrams showing, for each of 20 independent MONET training seeds, how per-seed cluster assignments (Sn:C0-C3) map onto the final consensus subtype assignment (Final:1-4) after seed-to-consensus alignment. Ribbon width is proportional to the number of donors following that path; ribbon color indicates the destination consensus subtype.

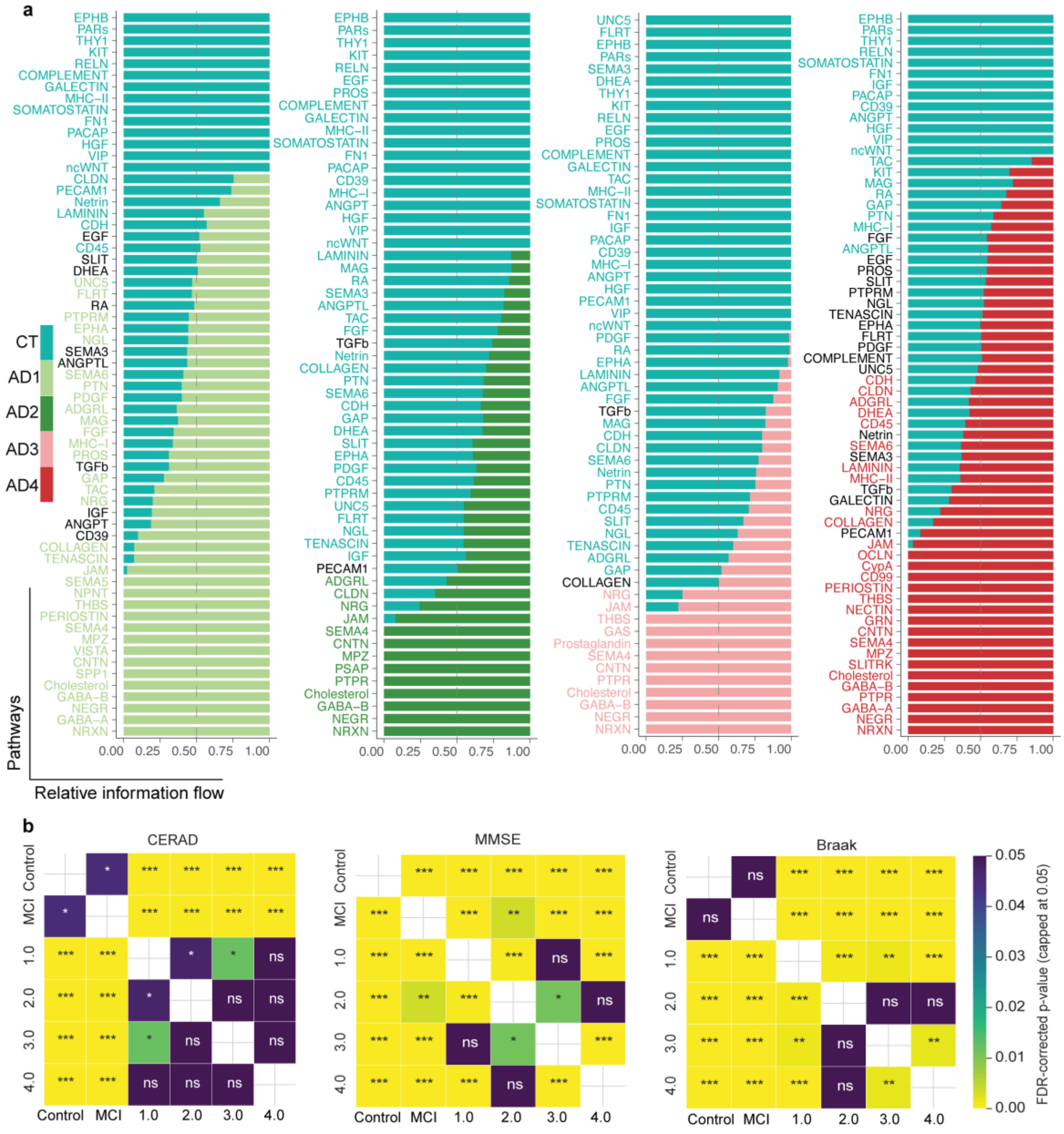


**Supplementary Figure 6 |** Subtype-specific cell-cell communication and clinicopathological association. **a,** Relative information flow of each significant signaling pathway across cognitively normal controls (CT) and the four molecular AD subtypes (AD1-AD4, color-coded). **b,** Pairwise statistical significance (Mann-Whitney U test, Benjamini-Hochberg FDR-corrected) of CERAD neuritic plaque score, Mini-Mental State Examination (MMSE) score, and Braak neurofibrillary tangle stage (left to right) across cognitively normal controls (Control), MCI, and the four molecular AD subtypes (1.0-4.0). Cell color indicates FDR-corrected p-value (capped at 0.05); *P < 0.05, **P < 0.01, ***P < 0.001; ns, not significant.

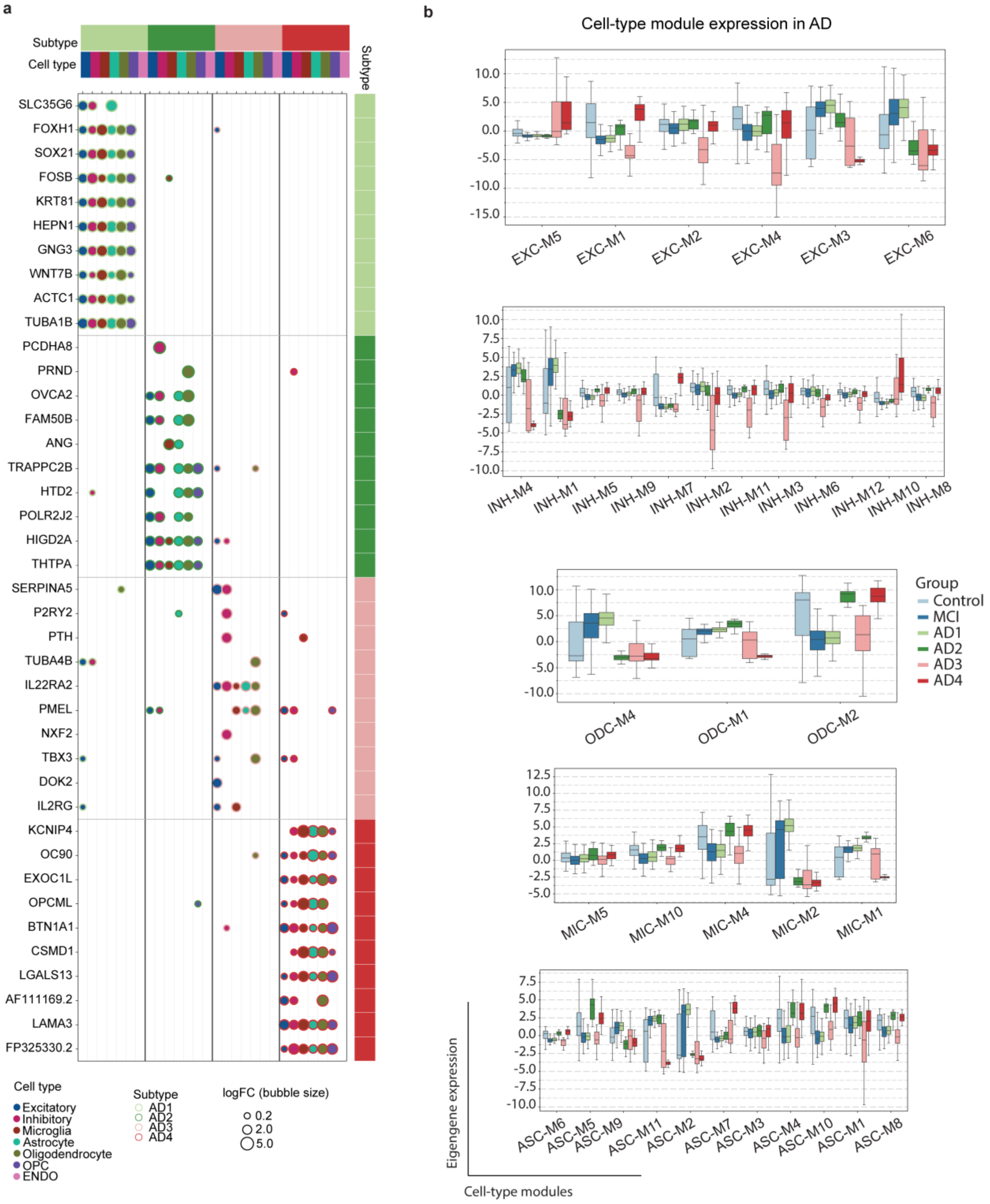


**Supplementary Figure 7 |** Subtype-specific marker gene expression and co-expression module activity. **a,** Balanced marker gene panel (10 genes per subtype, grouped by column) showing pseudobulk differential expression by cell type (bubble color) and magnitude (bubble size, log fold-change) across the four molecular AD subtypes. **b,** Eigengene expression of representative cell-type co-expression modules across cognitively normal controls (Control), MCI, and the four AD subtypes (AD1-AD4), grouped by cell type (excitatory neurons, inhibitory neurons, oligodendrocytes, microglia, astrocytes).

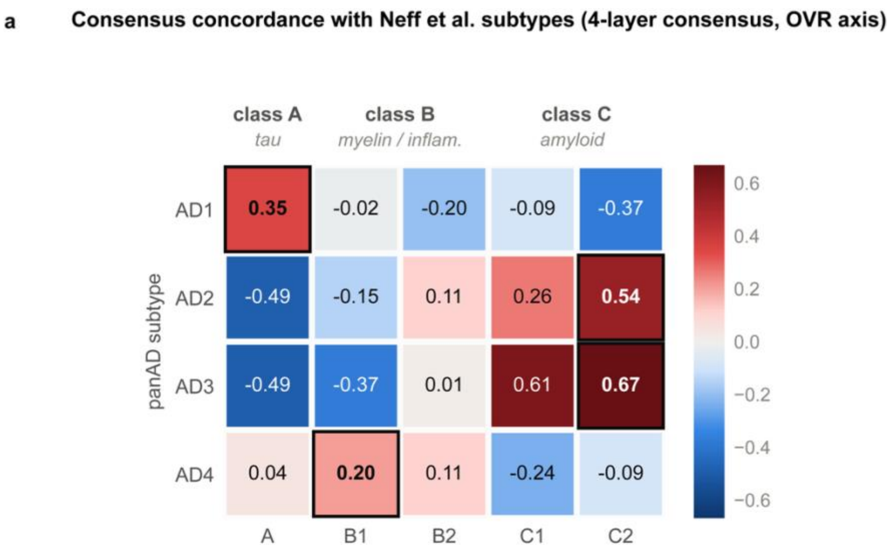


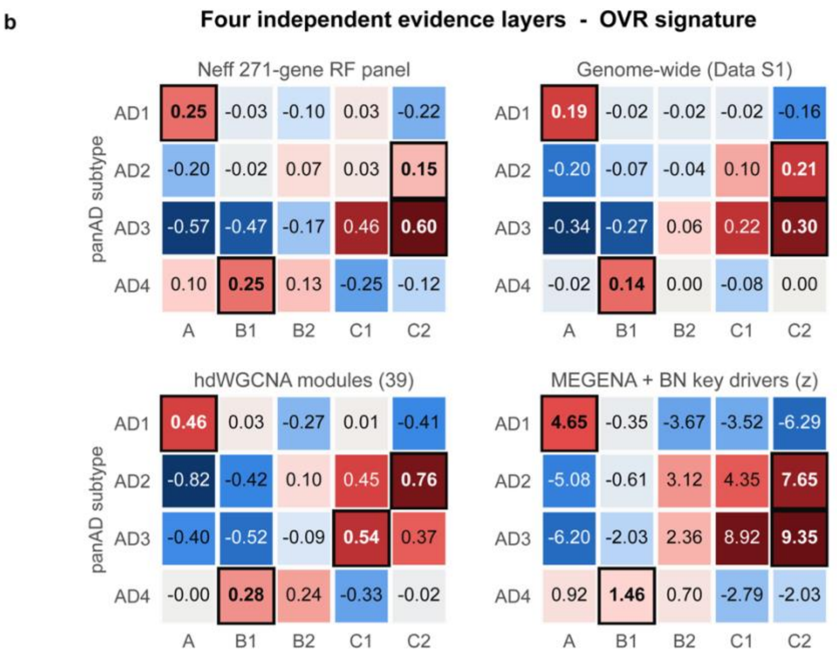


**Supplementary Figure 8 |** Reproducibility of panAD molecular AD subtypes in an independent cohort. **a,** Consensus concordance (Fisher-z-averaged correlation across the four independent evidence layers in b) between subtype-specific transcriptional signatures (one-vs-rest comparison) of the four panAD AD subtypes (AD1-AD4) and the five transcriptomic subtypes (A, B1, B2, C1,

C2; grouped into classes A/tau, B/myelin-inflammatory, and C/amyloid) defined by Neff et al. from bulk transcriptomic profiling of the independent MSBB cohort. Boxed cells indicate the best-matching Neff subtype for each panAD subtype (Hungarian assignment). **b,** The four independent evidence layers contributing to the consensus in a: the Neff 271-gene random-forest classifier panel, genome-wide correlation, hdWGCNA co-expression module scores, and MEGENA/Bayesian-network key-driver enrichment (z-score). Boxed cells indicate the best match per row within each layer. AD1 and AD3 showed concordant, statistically significant matches across all four evidence layers (class A and class C, respectively), whereas support for AD2 and AD4 was weaker and layer-dependent (see Methods).

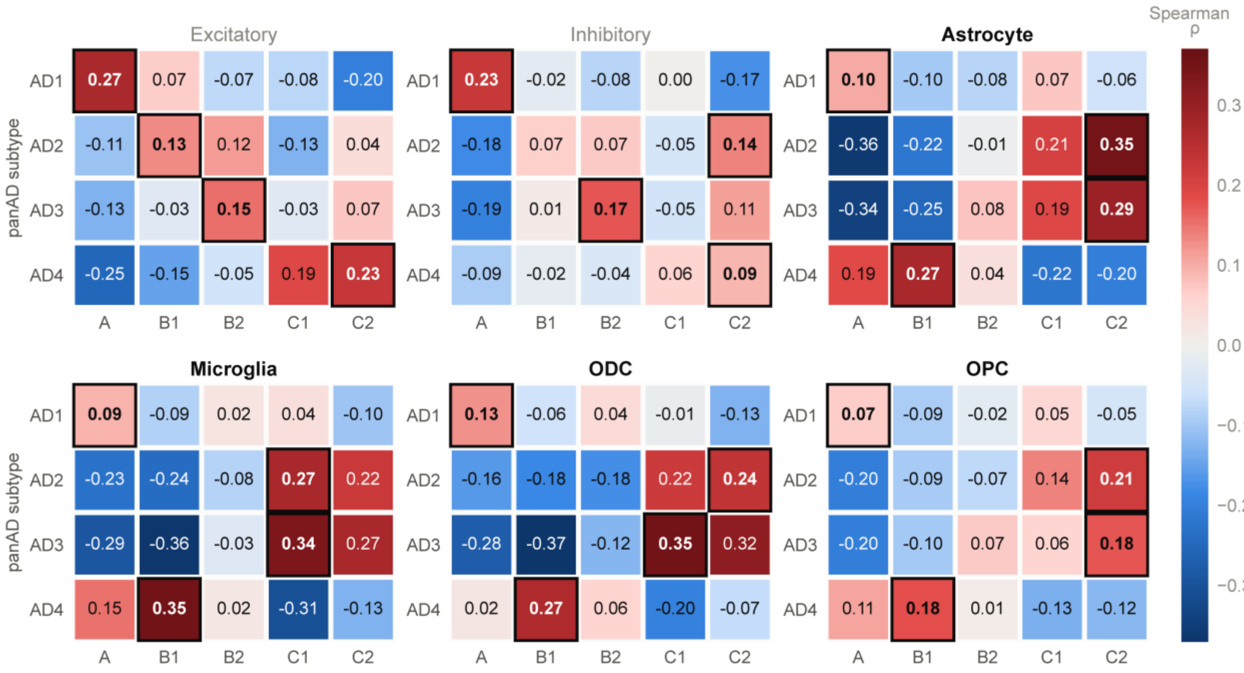


**Supplementary Figure 9 |** The panAD-Neff subtype concordance is primarily carried by glial cell types. Per-cell-type Spearman correlation between each panAD AD subtype (AD1-AD4) and the five Neff et al. MSBB subtypes (A, B1, B2, C1, C2), using the subtype-specific (one-vs-rest) signature, shown separately for excitatory neurons, inhibitory neurons, astrocytes, microglia, oligodendrocytes (ODC), and oligodendrocyte precursor cells (OPC). Boxed cells indicate the best-matching Neff subtype per row within each cell type. All four glial cell types independently recover concordant subtype matches, whereas both neuronal populations reproduce the AD1-class A match but show weaker or divergent correspondence for AD2-AD4, indicating that the panAD-Neff subtype concordance shown in Supplementary Fig. 8 is predominantly driven by glial transcriptional programs.
